# Empirical Validation of Composite Fractional Noise Models in Nanopore Signals

**DOI:** 10.64898/2026.09.28.754894

**Authors:** Dylan Charnock, Chalmers C. C. Chau, Paolo Actis, Christoph Wälti

## Abstract

Nanopore sensors have transformed single-molecule analysis, enabling real-time detection of biomolecules with unprecedented resolution. Understanding and modelling noise in nanopore sensing is essential to unlocking their full analytical potential. Although mechanistic models of nanopore noise exist, a rigorous statistical framework that characterises noise structure across diverse experimental conditions is still lacking. Here, we present a comprehensive empirical analysis of nanopore noise using twelve diverse experimental conditions acquired with different nanopores, analytes, and electronic recording systems. By employing multiple, mutually reinforcing statistical methods, we establish that nanopore noise is well described by a composite fractional model comprising multiple fractional Gaussian noise and fractional Brownian motion components. This finding is supported by confirmation of Gaussianity, rigorous interpretation of second-order exponents, multifractal analysis, and evidence that transient deviations are attributable to deterministic or experimental artefacts rather than alternative stochastic mechanisms. We further resolve quasi-deterministic structures, such as baseline trends and structural breaks, and demonstrate their separability from the underlying stochastic profile. Collectively, these results provide the first formal statistical validation of a composite fractional model for nanopore noise and establish a principled basis for noise reconstruction, realistic simulation, and statistically grounded algorithm design for nanopore sensing applications. This study provides the foundation for the creation of high-fidelity performance evaluation datasets for signal processing, and the controlled generation and augmentation of data for machine learning while maintaining statistical fidelity to experimental conditions.

## I. INTRODUCTION

Nanopore technology has entered the post genomics era. The sequencing of nucleic acids with nanopore is now routine and the field has now moved towards protein analysis and fingerprinting [1], [2], [3], [4], [5], [6], [7], [8], [9], [10], [11].

In a typical configuration, a nanopore separates two fluidic reservoirs filled with an electrolyte, each fitted with an electrode. The application of a fixed voltage generates a constant ion current which is stabilised by the balance between electrostatic drift, which drives ions along the electric field, and diffusion, which promotes uniform ion distribution. The nanopore constriction, the point of highest resistance, creates a highly concentrated electric field known as the capture region. In solution, charged analytes primarily undergo Brownian motion. However, upon entering the capture region, increasing electrostatic and electro-osmotic forces start to dominate and drive the analytes through the nanopore. These translocations of single molecules temporarily disrupt the ion current giving raise to measurable single molecule events. The ability to analyse single molecules in solution with high sensitivity, real-time resolution, and at low cost, has enabled a wide range of applications for nanopore sensing.

Nanopore noise is well studied mechanistically and plays a critical role in analysis [12], [13], [14], [15], [16], [17], [18], [19]. It determines the signal-to-noise ratio (SNR) and thus the limit of detection [20], which remains a critical challenge in small-analyte recordings. It also shapes event interpretation, since long-range dependent noise at the event scale distorts small-scale features, biases event statistics, and introduces uncertainty in baseline determination and, by extension, event bounding [21]. Most event extraction algorithms remove deterministic trends before applying statistical significance tests to the residual background noise [22], [23], [24], [25], [26], or detect first-order changes using thresholds derived from higher-order noise statistics [27], [28], [29]. In practice, denoising typically precedes extraction, together acting as a band-pass filter around event scales to suppress noise and improve algorithmic performance and interpretation. Taken together, noise is the central factor governing the applicability and performance of analysis pipelines, and it must be rigorously understood to maximise analytical accuracy.

The currently accepted model describes nanopore noise as a superposition of frequency-dependent regimes [13-14], with the Power Spectral Density (PSD) serving as the primary analytical tool. Nonlinearities in the low-frequency PSD have been observed in the presence of poly-ethylene glycol (PEG) and attributed to absorption kinetics [30], while theoretical limits to the power-law scaling have been proposed and validated through simulation and experimentation [31]. Gaussianity is often assumed to support statistical inference and to generate synthetic noise for machine learning or performance evaluation [32-34], but, to the best of our knowledge, this assumption has never been rigorously tested. Beyond stochastic properties, structural breaks have been linked to crowding effects [35], and stationary baseline components have been constructed using conductance models [36].

Despite extensive mechanistic studies, the statistical properties of nanopore noise remain largely unexplored, and a comprehensive statistical analysis has not yet been performed.

Such analysis is necessary to validate the interpretation of nanopore noise in a statistical context and to develop a representative model, which would support further characterisation of nanopore noise, the development of analysis algorithms, and the generation of synthetic data for signal processing and machine learning applications.

Here, we propose a data-driven approach to noise modelling, where assumptions are derived empirically from statistical analysis of a diverse set of nanopore traces. Rather than relying on predefined and physically motivated models, we apply a sequential series of tests to systematically build and validate assumptions, leading to the construction of a representative statistical model. These tests assess distributional properties, structural irregularities, dependence, second-order self-similarity, and multifractality. At each stage, individual divergences are identified, and multiple complementary methods are used to assess each property, enabling cross-validation between methods to reinforce conclusions. We do not attempt to model the non-stationary baseline component or low-frequency nonlinearities that occur on timescales far exceeding the event scale, but we aim to identify where such effects occur and how they influence higher-order fluctuation dynamics.

By identifying common characteristics across multiple noise profiles, we construct a generalised statistical model that captures the underlying noise structure irrespective of experimental conditions. This study provides the first formal statistical validation of a composite fractional model for nanopore noise, enabling principled reconstruction, realistic simulation, and statistically grounded algorithm design.

## II. METHODS

### II A. Data Collection

A total of twelve traces were analysed, with two acquired specifically for this study and ten obtained from public data repositories and research groups. A detailed summary of all traces, including specific conditions for each experiment, is provided in the Supplemental Material (Table SI). Data were collected under three different conditions: cis-trans translocation, trans-cis translocation [37], and single-cell nanoinjection [38]. Three amplifiers were used: Molecular Devices MultiClamp 700b (100 kHz), Elements Nanopore Reader 10 MHz (1 MHz), and Elements eOneB (20 kHz).

The analytes studied included 7 kbp double-stranded (ds)DNA, 3 kbp dsDNA, *λ* dsDNA, 12 nm nanostar [5], AAV9, 30 nm AgNP [37], DNA origami (“2×2”) [39], β-galactosidase, and 70S ribosome (New England Biolabs). Experiments were conducted using three types of nanopores: quartz nanopores (~20–160 nm diameter), SiN fabricated via controlled dielectric breakdown (~15 nm diameter) [40], [41], and Norcada NXPR5001Y-75 nm-AO-HR 20 nm SiNx + 60 nm SiO□ (~70 nm diameter).

Experiments were performed at voltages ranging from −700 mV to 300 mV, with analyte concentrations between 0.2 nM and 1 µM. The electrolytes used ranged from 100 mM to 1 M KCl, with one trace recorded in 10 mM KCl + 30 mM NaCl and another in 137.9 mM NaCl + 2.67 mM KCl. Measurements were conducted both with and without 25% or 50% (w/v) PEG 35k.

### II B. Event Removal

Event detection was performed using the iterative baseline method [23] combined with a two-tailed z-score test to set the threshold at the single expected outlier magnitude. Baseline cutoff frequencies were maximised while avoiding attenuation of the largest events, and the procedure was applied over two iterative passes to further refine the baseline estimate and the detection threshold. After both passes were completed, detected events were manually classified according to their waveform characteristics and their consistency with other detected events. Clear events were extracted, while potential artifacts or ambiguous events were retained.

A secondary visual windowed pass was applied using the same criteria, where traces were examined in non-overlapping segments of adjustable size. The default window size was 0.1s. Event boundaries in both initial detection and visual validation were determined algorithmically, using the first and last baseline crossings, and no manual adjustments were made for long-tailed events. Extracted events were removed entirely, and the remaining segments were concatenated directly at these boundaries. While this approach introduces some bias, it was preferred over interpolation or the introduction of synthetic segments, as those alternatives would themselves alter the signal and confound the subsequent noise analysis.

### II C. Visual Analysis

Visual inspection of the traces was used to generate initial statistical assumptions and identify prominent structures. These observations guided the selection and applicability of subsequent statistical tests, along with the expected model.

Each trace was plotted alongside the events-removed counterpart, stacked vertically, analogous to Supplemental Material (Fig. S1). Traces were examined at multiple scales using the previously described windowed pass method (Sec. II B). This multiscale inspection allowed for the identification of both global and local structures.

### II D. Quantile-Quantile and Probability-Probability

Quantile-quantile (Q-Q) and probability-probability (P-P) plots were computed to assess deviations from normality in each events-removed trace, with the Q-Q plot emphasising differences in the tails of the distribution and the P-P plot highlighting deviations in the centre.

Nonlinear detrending was applied to the traces using a cubic spline with an equivalent 1 Hz cutoff frequency, ensuring the removal of quasi-deterministic trends without affecting the residual distribution across all event scales. The sample data were fitted to both normal and skew-normal comparison distributions using maximum likelihood estimation (MLE). Quantiles and cumulative distribution functions (CDFs) were computed for each sample and both theoretical distributions.

The goodness of fit was assessed by calculating the coefficient of determination (R^2^) and the mean relative error (MRE) between each sample and the theoretical quantiles/CDFs. Additionally, the standard error of the sample quantiles/CDFs was determined to construct a 95% confidence interval around each reference line.

### II E. Complementary Cumulative Distribution Function

Complementary cumulative distribution functions (CCDFs) were computed to assess deviations in distribution tails relative to the normal distribution for the traces with the events removed.

Nonlinear detrending was applied to the traces using the previously described method (see Sec. II D). Each sample was fitted to the theoretical normal distribution using MLE. To analyse tail behaviour, each sample was split at the mean, with the upper and lower portions representing conductive and resistive tails. The theoretical distributions were similarly divided to define reference tails.

CCDFs were computed for the samples and theoretical distributions independently and for each tail. The goodness of fit was evaluated by calculating the R^2^ score and the MRE between sample tails and the theoretical references.

### II F. Distributional Point Estimates

To characterise the distributional properties of the events-removed traces, several point estimates were computed after detrending using the previously described method (see Sec. II D).

The Shapiro-Wilk *W* statistic [42] was computed as a point estimate of departure from normality. The Fisher-Pearson coefficient of skewness quantified the asymmetry of the sample distributions, while the Fisher coefficient of kurtosis measured the tendency for extreme values to occur relative to normal distributions. Here, skewness and kurtosis refer to the standardised third central moment and the excess standardised fourth central moment, respectively, while the Shapiro-Wilk *W* statistic is conventionally used to test the null hypothesis that a sample is drawn from a normal distribution.

### II G. Rescaled Range

The rescaled range (R/S) method [43], [44] was applied to evaluate statistical dependence within the events-removed traces, assuming stationarity.

For each trace, log-spaced scales were selected, ranging from one-eighth of the total trace length down to a numerically stable lower bound (~2^4^). For each scale, n, the trace was divided into non-overlapping segments of length n. Within each segment, the mean was computed, followed by the cumulative deviation from the mean. The adjusted range *R*_*i*_(*n*) was determined as the difference between the maximum and minimum of the cumulative deviation. The standard deviation *S*_*i*_ (*n*) was computed for each segment, and the rescaled range *R*/*S*(*n*) was obtained as the average of R_i_(n)/S_i_(n) across all segments. Finally, *log*(*n*) was plotted against *log*(*F*_*n*_), where *F*_*n*_ represents the mean *R*/*S* ratio at scale *n*. The Hurst exponent *H* was estimated as the slope of this relationship using least squares regression.

### II H. Detrended Fluctuation Analysis

Detrended Fluctuation Analysis (DFA) [45] was applied to evaluate statistical dependence within the events-removed traces without assuming stationarity. This method generalises the Hurst exponent *H* in terms of *α*, allowing *α* to exceed one, thereby capturing long-range dependence in both stationary (*α* = *H*) and non-stationary (*α* = *H* + 1) time-series (see Tables SII and SIII).

For each trace, log-spaced scales were selected, ranging from one-eighth of the total trace length down to a numerically stable lower bound (~2^4^), resulting in 40 logarithmically spaced scales. This selection ensures statistical robustness based on extrapolated empirical confidence intervals from [46] and the minimum trace length used. DFA computations were performed using the Fathon implementation [47] with polynomial detrending orders *N* ranging from one to four. The optimal detrending orders were determined following the approach in [48], based on changes in slope and scaling regime crossovers in the *log*(*F*_*n*_) vs. *log*(*n*) plots.

After selecting the optimal detrending orders, *S* scaling regimes were identified per trace by fitting piecewise linear models to the *log*(*F*_*n*_) vs. *log*(*n*) plots. The number of regimes and their crossover points were determined by minimising the mean squared error (MSE) of a piecewise fit, optimised using the Nelder-Mead algorithm. For each identified scaling regime, the *α* parameter was extracted as the slope of the corresponding linear segment in the *log*(*F*_*n*_) vs. *log*(*n*) plot. The goodness of fit for each scaling regime was assessed using the R^2^ score.

### II I. Power Spectral Density

The power spectral density (PSD) was estimated to assess spectral scaling properties of both the raw traces and the traces with the events removed, specifically focusing on enhancing high-frequency resolution and enabling comparison with the time-domain scaling methods.

PSDs were computed using Welch’s method with a Hann window function to minimise spectral leakage, improve frequency resolution, and reduce discontinuities at segment boundaries. A 50% window overlap was applied to ensure statistical stability and sufficient spectral averaging, therefore maximising the number of independent segments contributing to each PSD estimate.

Window sizes were determined systematically based on the autocorrelation function (ACF) of each events-removed trace, ensuring that each window encompassed at least twice the lags at which the ACF fell and remained below 0.1.

### II J. Scaled Stationarity

The Kwiatkowski-Phillips-Schmidt-Shin (KPSS) [49] and the Augmented Dickey-Fuller (ADF) tests [50] were applied to assess the stationarity and ensemble dependence properties of the traces with the events removed. These tests were conducted on both the traces and a reference set of fractional long-range and short-range dependent time-series (0< *α* < 1.5) generated with one million samples using the Davis-Harte method [51].

The KPSS test evaluates the null hypothesis *H*_O_ that the series is trend stationary, with the alternative hypothesis *H*_l_ indicating non-stationarity. It has been shown to be consistent against both stationary and non-stationary long-range dependent alternatives (*I* (*d*), *d* > 0) [52]. Here, *I* (*d*) denotes a process integrated of order *d*, where *d* parameterises the degree of fractional integration (see Tables SII and SIII).

The ADF test evaluates the null hypothesis *H*_O_ that the series contains a unit root (*I*(d), *d* = 1), with the alternative hypothesis *H*_l_ suggesting that no unit root is present (*I* (*d*), *d* ≠ 1). This test has been shown to reject consistently the null for fractionally integrated series [53].

To evaluate temporal changes in stationarity and dependence, each trace was divided into non-overlapping segments of length n, corresponding to a 1 Hz period. This segmentation ensured high statistical power for both tests while enabling an assessment of temporal evolution across multiple segments. The KPSS test statistics were computed using the Newey-West heteroskedasticity and autocorrelation consistent (HAC) estimator to account for serial correlation, with the lag lengths selected using the data-dependent method [54]. The ADF test statistics were computed using Akaike Information Criterion (AIC) to determine the optimal lag lengths. For both tests, critical values at *α* = 0.01 were obtained from precomputed Monte Carlo simulations.

The distribution of test statistics across segments was computed and compared to those of the simulated time-series to assess how the stationarity and dependence properties of each trace compared to the reference set. The Earth Mover’s Distance (EMD) was used to identify the closest matching reference distributions via the Wasserstein metric. Additionally, the temporal evolution of the test statistics and results were visualised alongside the respective trace to identify changes in dependence properties and potential intermittence.

#### II J 1. Scaled Stationary Interpretation

Together, the aforementioned tests provide complementary insights into stationarity and dependence properties. When *d* is close to zero, both tests indicate trend stationary. Conversely, when *d* is close to one, both tests suggest non-stationary. If the KPSS test rejects stationarity while the ADF test fails to reject the unit root, the series is likely fractionally integrated [52], [53], [54].

#### II J 2. Scaled Stationary Novelty Statement

To our knowledge, the combined segmented application of KPSS and ADF tests, applied to high-resolution non-overlapping segments and compared to a reference set of fractional time-series, represents a novel approach to assessing time-local stationarity and dependence properties in stochastic traces. This methodology allows for detailed characterisation of dynamic changes in stationarity and dependence behavior, which are not accessible through conventional global stationarity tests, and provides a rigorous basis for comparing experimental noise profiles to fractional noise models.

### II K. Multifractal Detrended Fluctuation Analysis

Multifractal Detrended Fluctuation Analysis (MF-DFA) [55] was applied to evaluate fluctuation-dependent statistical properties of the events-removed traces and their shuffled counterparts, which emerge when small and large fluctuations exhibit different scaling behaviours. The method extends standard DFA by introducing a range of moment orders, denoted by the parameter *q*, which allows for differential weighting of small and large fluctuations. Here, DFA *α* is equivalent to MF-DFA *α*_2_, corresponding to the second-order fluctuation scaling. By varying *q*, one can assess how fluctuations of different magnitudes contribute to the overall scaling behaviour, with positive *q* emphasising larger fluctuations and negative *q* emphasising smaller fluctuations.

For each trace, log-spaced scales were selected, ranging from one-eighth of the total trace length down to a numerically stable lower bound (~2^4^), resulting in 40 logarithmically spaced scales. MF-DFA computations were performed using the Fathon implementation [47], applying the same polynomial detrending orders previously determined from *N*-DFA (see Sec. II H). The analysis incorporated eleven *q*-orders, ranging from −4 to 4, to capture both small and large magnitude fluctuations.

For each trace and *q*-order, *S* scaling regimes were fitted using the previously determined crossover points and piecewise fitting methodology to obtain the *q*-order scaling exponents *α* _*q*_ (see Sec. II H). Each *α* _*q*_ was then transformed into the *q*-order mass exponent *τ*_*q*_ using the standard relation:

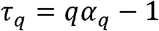

The Hölder exponent *h*_*q*_ was then computed as the discrete derivative of *τ*_*q*_ with respect to *q*. This notation is adopted here for consistency with DFA-based notation used throughout:

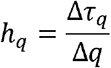

Finally, the multifractal spectrum *D*_*q*_, describing the distribution of singularity strengths across fluctuation magnitudes, was computed using:

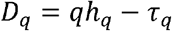

#### II K 1. Multifractal Detrended Fluctuation Analysis Interpretation

MF-DFA provides a powerful tool for characterising heterogeneous scaling behaviour, intermittency, and distributional structure in time-series data. Its strength lies in its ability to disentangle complex fluctuation dynamics across both temporal scales and fluctuation magnitudes, enabling systematic classification of scaling regimes and their underlying mechanisms [55], [56]. This is particularly valuable in the context of nanopore data, where manual identification of millions of samples is not feasible.

Nonlinear interactions can introduce fluctuation-dependent correlations, leading to variations in the scaling exponent across fluctuation magnitudes and producing or modifying vertical separation between *q***-**orders in the *log*(*F*_*n*_) vs. *log*(*n*) plot. Localised structural breaks appear as discontinuities as they do not affect all scales and *q*-orders uniformly. In contrast, genuine multifractality manifests as stochastic multiscale irregularity or fluctuation-dependent changes in scaling behaviour. Variance changes may remain undetectable, induce a gradual increase in the scaling regime slope, or produce fan-like spreading in which higher *q*-orders diverge. Additionally, *q*-order slopes may remain unchanged while exhibiting vertical separation if multiple noise sources alter the fluctuation distribution, indicating shifts in the relative weighting of fluctuation magnitudes without a corresponding change in the scaling exponent.

To determine whether variance scaling originates from temporal correlations or distributional effects, a random shuffling procedure can be applied, preserving the marginal distribution while eliminating temporal correlations. If vertical separation between *q*-orders collapses and the scaling behaviour transitions to that of uncorrelated white noise post-shuffling, this indicates that both multifractality and scaling are driven by temporal correlations. Conversely, if vertical separation or correlated scaling behaviour persists, this suggests that distributional features such as heavy tails, variance shifts, or mixed noise sources contribute to the observed multifractality and scaling.

Intermittency or correlation structures that are not apparent in the PSD or standard DFA can be revealed through MF-DFA, which captures higher-order fluctuation dynamics beyond the scope of conventional second-order analyses.

### II L. Time-dependent Detrended Fluctuation Analysis

Time-dependent Detrended Fluctuation Analysis (T-DFA) [56] was applied to investigate multifractality at specific scales in the events-removed traces. Unlike MF-DFA, T-DFA focuses on the temporal evolution of individual scales relative to the maximum scale considered. Instead of a non-overlapping segmentation approach, T-DFA employs a sliding window to maximise time resolution, enabling a continuous assessment of how scaling properties evolve within a given scale. The resulting local scaling exponent *α*_*t*_ represents the local DFA scaling exponent estimated at time *t*, thereby providing a time-resolved view of multifractality.

For pertinent traces, a subset of scales exhibiting multifractal effects was selected for analysis. T-DFA computations were performed using the Fathon implementation [47], with the local scaling exponents *α*_*t*_ computed relative to the associated scaling regime determined by DFA, and using the same detrending order established via *N*-DFA (see Sec. II H). The resulting *α*_*t*_ were then plotted alongside the corresponding traces to illustrate temporal dynamics and identify the origins of multifractality.

#### II L 1. Time-dependent Detrended Fluctuation Analysis Interpretation

In relation to MF-DFA, *q*-order separation can be inferred from the variance of the local scaling exponent *α*_*t*_ within a chosen scale: low *α*_*t*_ correspond to large *q*-orders (i.e., decrease in relative slope), whereas high *α*_*t*_ reflect low *q*-orders. Unlike MF-DFA, which aggregates scaling behaviour across the entire trace, T-DFA enables temporal localisation of changes in scaling structure. By tracking these fluctuations over time, T-DFA provides direct insight into transient irregularities such as structural breaks, intermittency, and scale-dependent variations within the signal. This scale-specific analysis enables direct interpretation of transient features and whether their origin is intrinsic to the system, thereby informing their inclusion in the resultant noise model.

## III. RESULTS AND DISCUSSION

To ensure a representative noise characterisation, we analysed twelve solid-state nanopore traces, recorded by six researchers from three independent laboratories under different experimental conditions, including a range of biochemical buffers and analytes, using three different amplifiers, with nanopores fabricates using different methods. We use a combination of nanopore traces recorded with (ten traces) and without (two traces) an analyte as analyte interaction heavily influence the baseline and noise environment through charge density modulation, Coulomb interaction, and potential absorption or blocking mechanisms [35], making a statistical understanding of these effects essential for accurate modelling. For an overview of each trace, see Sec. II A, Fig. S1, and Table SI. For event removal details, see Sec. II B, and Fig. 1(a).

**Fig. 1.**
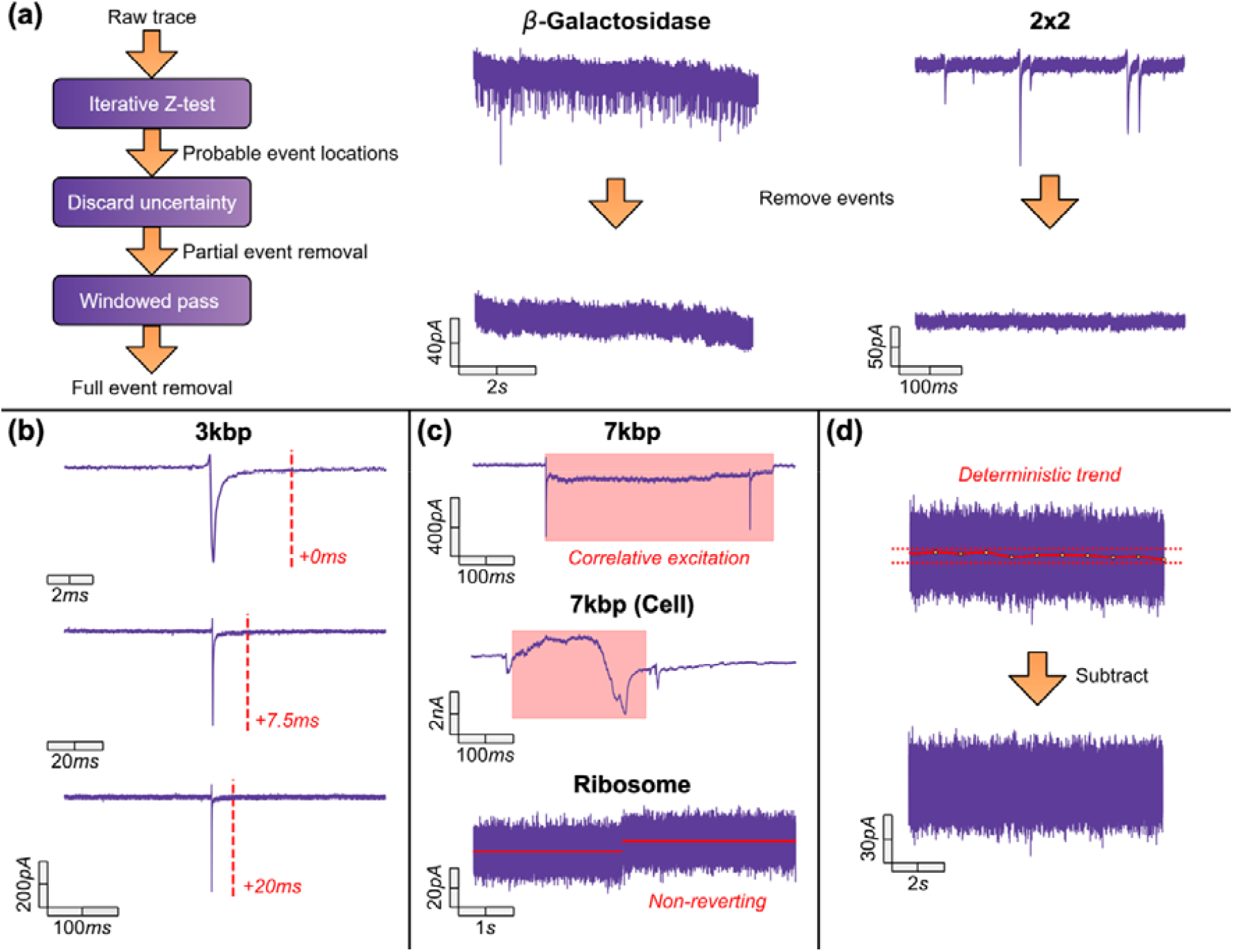
(a) Flow chart showing the event removal process and examples at coarse and fine scales for ion current traces of two datasets (β-galactosidase and 2×2 DNA origami); (b) Zoomed section of a single event showing asymptotic tails in the 3 kbp DNA trace demonstrating scale-free behaviour, illustrating that the endpoint (dotted red lines) would be detected at different positions based on the zoom factor. The stated times are the differences between each endpoint and the endpoint determined in the top example; (c) Examples of the three types of structural breaks observed in the datasets. The red boxes highlight reverting breaks, and the red lines represent the two distinct conductance levels for a non-reverting break; and (d) a representative example of deterministic trend removal. The red line with the orange dots shows the extracted trend, and the dotted parallel lines aid in interpretability. Corresponding full trace views are shown in Fig. S1.

The first step involves a visual inspection of the traces to generate initial assumptions and identify prominent structures. These observations guide the selection and applicability of subsequent statistical tests. Each trace, with events removed, is plotted alongside the raw trace and analysed through multiscale inspection, enabling identification of both global and local structures (see Sec. II C).

Visual inspection reveals that most of the traces exhibit deterministic trend stationarity, with no apparent structural breaks. The Ribosome trace contains a single, non-reverting structural break in the mean, characterised by a sharp transition into a less conductive state and a preservation of higher-order distributional properties (Fig. 1(c)). No changes are observed beyond the first order, and the statistical properties before and after the break appear identical. This behaviour likely reflects irreversible analyte absorption or a permanent alteration to the system, independent of stochastic effects. The 7 kbp and 7 kbp (Cell) traces exhibit frequent and sharp structural breaks in the mean, which are often preceded by an event and always induce a temporary increase in serial correlation (Fig. 1(c)). These breaks always revert, either gradually, or by a similarly sharp break after a temporary change in conductance. However, reversion does not always result in the original conductance state observed before the break. Importantly, structural breaks were found to be associated with an increased translocation event frequency, larger resistive events, occur more frequently following the minimum conductance of each trace and always transition to a higher conductance that subsequently reverts to a lower conductance. This behaviour is indicative of blocking mechanisms induced by analyte interaction with the pore. The sharp increases are characteristic of desorption, unblocking or rearrangement and the dual reversion mechanisms may reflect reaggregation or reabsorption. Similar behaviour has recently been reported and attributed to analyte crowding in the nanopore, although distinct due to the polarity of the relative conductance states [35].

As mentioned, a subset of traces recorded in the presence of PEG display post-translocation asymptotic tails, manifesting at very low frequencies compared to the event scale. These tails complicate event endpoint identification in otherwise stationary environments, as their interpretation is scale-dependent (Fig. 1(b)). Notably, these tails may overlap with nonlinear baseline fluctuations, potentially introducing bias in subsequent analyses.

In reference to distributional properties, the variance of most traces appears constant across time and scales, with no significant heteroskedasticity observed. When structural breaks are excluded, the 7 kbp and 7 kbp (Cell) traces exhibit similar behaviour. Additionally, noise distributions appear symmetric, with no evidence of heavy tails across all scales examined.

In essence, most traces can be characterised as stationary around a deterministic trend, which manifests only at low frequencies. Although the 7 kbp and 7 kbp (Cell) deviate from this characterisation due to structural breaks, when these breaks are excluded, their behaviour aligns with the rest of the traces (Fig. S1). Therefore, for subsequent analysis, we proceed under the assumption of deterministic trend stationarity. That is, removal of deterministic trends yields an approximately stationary series.

Assessing the distribution of each noise profile is essential for subsequent statistical analyses. First, both the KPSS [49] and ADF [50] tests rely on assumptions consistent with normally distributed or finite-variance residuals (see Sec. II J). In addition, the interpretation of DFA [45] depends on the distributional properties of the series, as the scaling behaviour of partial sums is directly linked to the rate of variance growth, which is influenced by the underlying distribution (see Sec. II H). Similarly, PSD and autocorrelation function (ACF) analyses rely on assumptions of finite variance and are particularly sensitive to heavy-tailed distributions (see Sec. II I). In the presence of heavy tails, these methods will produce misleading indications of long-range dependence, as extreme events dominate the variance and distort correlation structures. Specifically, heavy-tailed distributions, such as alpha-stable processes with stability parameter *α* < 2, can induce artificially slow-decaying autocorrelations and spurious *f*^-−*γ*^ scaling in the PSD, even when no genuine long-range dependence is present.

Beyond these considerations, distributional testing also allows us to assess whether the structural breaks observed in the 7 kbp and 7 kbp (Cell) traces are associated with an underlying distribution distinct from the rest of the test set, and to assess whether the conventional assumption of Gaussianity remains valid upon removal of the deterministic trend.

Prior to distributional assessment, a nonlinear detrending at 1 Hz was applied to each trace to remove low-frequency baseline fluctuations while preserving event scales of interest. This was done in accordance with our pre-established assumptions and on the events-removed traces (Fig. S1). Additionally, to assess whether their steady-state distributional properties were consistent with those observed in the remaining traces, the 7 kbp and 7 kbp (Cell) traces were truncated to segments of at least one million samples that avoided areas with structural breaks. These sub-traces were analysed separately, alongside the original full traces, in all subsequent analyses.

Three complementary approaches were employed to characterise the distributional properties. First, Q-Q and P-P plots were generated with 95% confidence intervals and R^2^ fit measures against both normal and skew-normal distributions (see Sec. II D, Figs. S2-S5). Q-Q plots emphasise tail behavior, while P-P plots focus on central distribution alignment. Second, CCDFs were used to examine conductive and resistive tails when compared to normal distributions (see Sec. II E, Fig. S7). Finally, point estimates of skewness, kurtosis, and overall deviation from normality were computed using the Fisher-Pearson coefficient of skewness, Fisher coefficient of kurtosis, and Shapiro-Wilk *W* statistic [42], respectively (see Sec. II F). Together, these approaches provided a comprehensive assessment of distributional properties relevant to future analyses and characterisation of time-localised structures.

Analysis of the Q-Q and P-P plots indicates that most traces exhibit distributional properties consistent with normality. Specifically, ten out of twelve traces fall within the 95% confidence intervals in the P-P plots, demonstrating strong agreement with normal distributions in the central regions. Of these, eight traces also remain within the 95% confidence intervals in the Q-Q plots, further supporting normality with respect to tail behavior (Figs. S2-S3). Within this subset, the two traces recorded without an analyte show substantial agreement with normality and do not exhibit contrasting behavior to the traces recorded with an analyte. Notably, the AAV9 trace is in near-perfect agreement with the normal distribution (Fig. 2(c)).

**Fig. 2.**
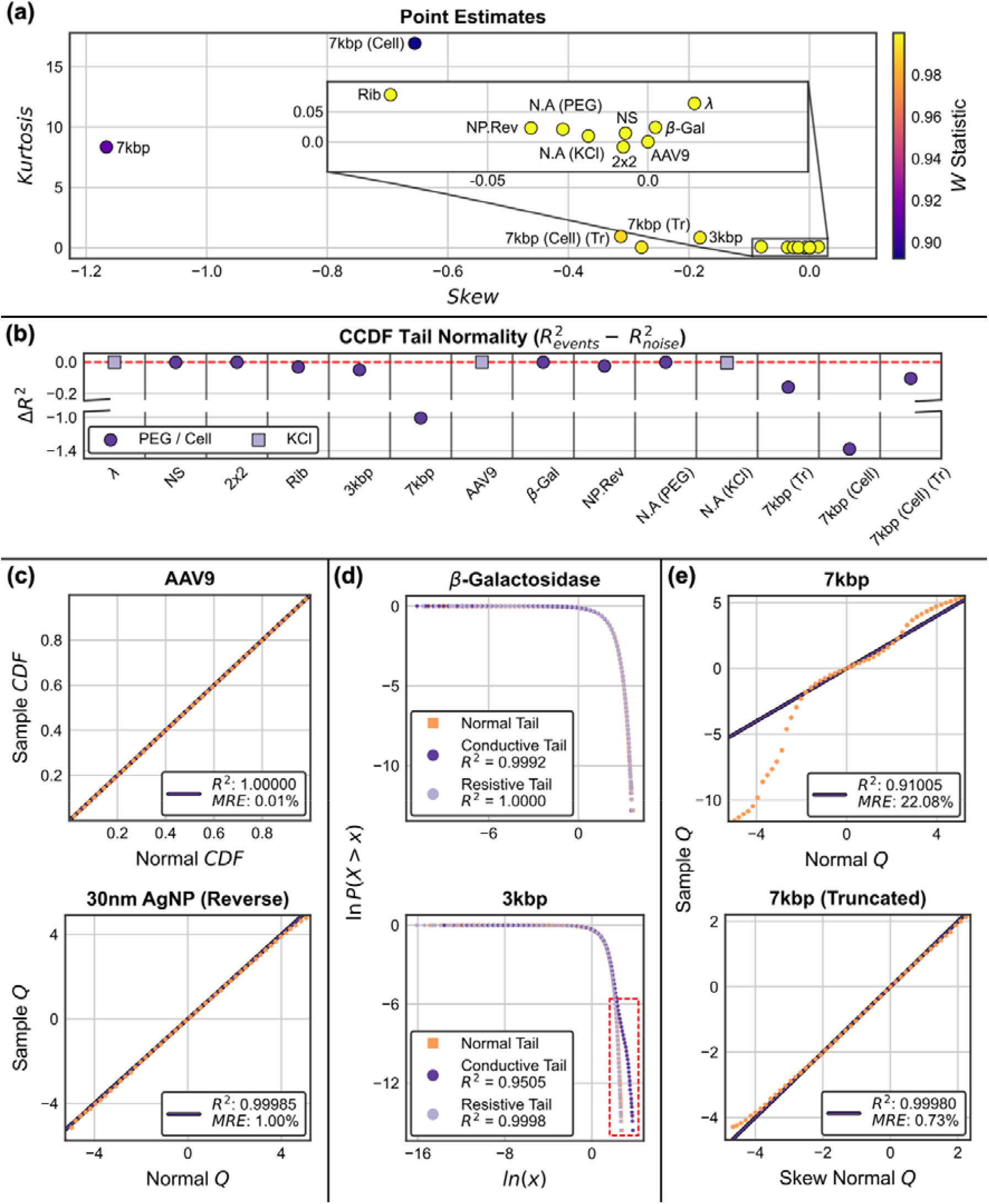
(a) Distributional point estimate results for each trace, the inset shows a zoomed-in representation of the traces concentrated around zero skew and kurtosis; (b) ensemble CCDF results demonstrating distributional divergence localised to the event side, where the red line at zero represents no difference in normality statistic between event and noise tails; (c) examples of P-P and Q-Q results, showing close adherence to the purple lines, which represent a perfect fit; (d) average (-Galactosidase) and divergent (3 kbp) CCDF examples, displaying tails that overlap the normal reference tails (orange squares), and event tail divergence for the 3 kbp trace (red box); and (e) the 7 kbp trace before and after truncation, demonstrating reversion to behaviour in line with the other traces. Abbreviations: Nanostar – NS; Ribosome – Rib; ***β***-Galactosidase – ***β***-Gal; 30 nm AgNP (Reverse) – NP.Rev; No Analyte (PEG) – N.A (PEG); No Analyte (KCl) – N.A (KCl); 7 kbp (Truncated) – 7kbp (Tr); 7 kbp (Cell) (Truncated) – 7kbp (Cell) (Tr).

The remaining two traces display asymmetric divergence. First, the Ribosome trace exhibits a negative skew accompanied by a heavy conductive tail. Further analysis confirms that this skew arises from the single structural break, with additional contribution from the placement of a nearby spline breakpoint used in the detrending process. Comparison to a skew-normal distribution aligns the Ribosome trace with the distributional properties observed in the other traces (Figs. S4-S5). Second, the 3 kbp trace displays extreme conductive tail divergence, consistent with previous observations of severe asymptotic event tails in this trace. Other traces recorded with 50% (w/v) PEG 35K, including the Ribosome trace when accounting for skew, exhibit similar but less pronounced divergences. Importantly, such tail divergences consistently occur on the side where events were present, suggesting that they reflect event-related artifacts rather than intrinsic distributional asymmetry.

The final two traces, the 7 kbp and 7 kbp (Cell), show major deviations from normality, displaying both leptokurtic and skewed behavior. Upon truncation to avoid structural breaks, both traces begin to align with the remainder of the set, although some divergences persist. Interestingly, the truncated traces display a combination of effects observed in both the 3 kbp and Ribosome traces. As the 7 kbp trace was recorded under similar experimental conditions to the 3 kbp trace, and the 7 kbp (Cell) trace within a crowded intracellular environment, the emergence of asymptotic tails is to be expected. This effect is most apparent in the 7 kbp (Cell) trace, where the truncated segment contains a markedly higher density of event removal sites compared to the truncated segment of the 7 kbp trace. Moreover, as with the Ribosome trace, both 7 kbp and 7 kbp (Cell) retain prominent skewness following truncation, driven by smaller structural breaks that could not be avoided when truncating. Nevertheless, comparison with skew-normal distributions fully restores alignment with the distributional properties observed in the other traces, indicating that these deviations are primarily attributable to a combination of event-related and structural artifacts rather than fundamental differences in distribution (Fig. 2(e)). CCDF analysis corroborates the tail behavior observed in the Q-Q plots, confirming that divergence occurs exclusively in the tail associated with event direction, while the opposite tail remains in close agreement with a normal distribution (see Fig. 2(b) and Fig. 2(d), and Fig. S7). In the case of skewed traces, an apparent shift affecting the tail without events is visible in the CCDF plot, which arises from the influence of the mean calculation when splitting the trace for analysing the tails separately. Specifically, as the degree of skew increases, the mean is increasingly pulled toward the heavier event-associated tail, causing the opposite tail to appear artificially lighter. Thus, the extent of this shift directly reflects the magnitude of skew present in the trace and should be interpreted as an artifact of mean displacement rather than a true deviation from normality in the opposing tail. Nevertheless, statistical comparisons consistently identify the event-associated tail as the primary source of divergence.

As final confirmation, distribution fitting analyses show that no alternative distribution provides a better fit than either the normal or skew-normal in all cases. In particular, alpha-stable distributions with stability parameter *α* ≠ 2 are not supported, ruling out heavy-tailed alpha-stable processes as an appropriate model.

In summary, these results demonstrate that the noise profiles are well described by Gaussian distributions in all but two cases. While these two traces exhibit heavier tails and pronounced skewness, their deviations are explainable as a combination of effects observed in other traces and are best captured by skew-normal distributions rather than heavier-tailed alternatives, provided structural breaks are avoided. These findings further suggest that structural breaks are not inherent distributional features, but instead quasi-deterministic artefacts superimposed on otherwise Gaussian noise. Importantly, tail divergence is consistently localised on the side where events were removed, primarily reflecting asymptotic tails identified in prior analyses, whereas the opposing tail is well-aligned with normality.

This analysis confirms that a 1 Hz cubic spline or equivalent nonlinear subtraction is sufficient to remove prominent low-frequency baseline fluctuations. Moreover, these findings validate the use of Gaussian-based statistical methods in subsequent analyses, thereby supporting the assumptions required for tests of stationarity, and the interpretation of scaling and dependence structures that follow.

Assessing self-similarity and dependence is essential for characterising the stochastic properties of each noise profile and whether models such as fractional Gaussian noise (fGn) or fractional Brownian motion (fBm) [57] are appropriate. First, establishing second-order self-similarity verifies whether the system exhibits scale-free behaviour, as expected for processes governed by power-law variance scaling. Here we adopt the term in the broad Mandelbrot sense, encompassing both persistent and anti-persistent regimes. In addition, identifying multiple scaling regimes allows us to detect superpositions of fractional components, consistent with the expected dominance of different noise sources at specific frequencies. Moreover, determining whether the scaling exponents and observed regimes correspond to stationary or non-stationary behaviour is critical for interpreting dependence, variance scaling, and the underlying process. Finally, assessing deviations from ideal scaling enables the identification of nonlinearities, band-limited effects, or additive components that may distort variance scaling and arise from instrumental or system-specific factors rather than intrinsic stochastic properties. Alongside distribution analysis, this approach provides a coherent description of each noise profile’s stochastic properties within a fractional noise framework.

Four complementary methods were employed to characterise second-order self-similarity and dependence properties in the events-removed traces and truncated versions of the 7 kbp and 7 kbp (Cell) traces. For the cross-method exponent correspondences and the interpretation of the resulting scaling behaviour, see Tables SII and SIII. First, R/S [43], [44] analysis was performed under stationary assumptions to provide a singular estimate of the Hurst exponent (see Sec. II G, Fig. S8). Second, DFA [45] was used to quantify variance scaling and to estimate multiple scaling exponents without assuming stationarity, following optimal detrending orders determined via -DFA [48] and segment sizes extrapolated from empirical confidence intervals [46] (see Sec. II H, Figs. S9-S10). Third, PSD analysis was used to assess spectral scaling, focusing on the increased resolution at high-frequencies beyond the numerical accuracy of the time-domain methods (see Sec. II I, Fig. S11). Finally, KPSS [49] and ADF [50] tests were applied to 1 Hz segmented traces to evaluate stationarity and ensemble dependence properties, interpreted as measures of fractional integration, over time. These tests were also applied to a reference set of fractional long-range and short-range dependent time-series to examine whether properties aligned with fractional noise models (see Sec. II J, Figs. S12-S20). Together, these approaches provided a robust multi-aspect characterisation of self-similarity and dependence across all traces.

Analysis of the R/S results revealed that multiple scaling regimes are present in all traces except for the 7 kbp (Cell) trace, which exhibits a single dominant regime. Divergence at larger segment sizes indicates deterministic non-stationarities impacting scaling. Consistent with this, N-DFA identifies third-order detrending as optimal without exceptions, in agreement with the detrending order applied during distributional analysis (Fig. S9). After applying this detrending order, DFA resolves one or two well-defined scaling regimes in all *λ* traces, with negligible deviations from self-similarity in eight out of twelve cases (Fig. S10). Although a gradual reduction in the scaling exponent at the smallest scales is apparent in several traces. Among the divergences from self-similarity, the 7 kbp, No Analyte (PEG) and *λ* traces exhibit deviation associated with line-frequency or its first harmonic, while the No Analyte (KCl) trace shows a pronounced bump in the large-scale regime similar to features reported by Knowles et al. [30] in traces recorded with PEG (Fig. 3(c)). However, this cannot be attributed to PEG or residual events. Non-stationary regimes (*α*> 1) are observed only in the 7 kbp, 7 kbp (Cell), *λ*, and *β*-galactosidase traces. The 7 kbp, λ, and β-galactosidase traces transition to stationary behaviour at approximately 155 Hz, 3.2 Hz, and 8.4 Hz, respectively. The 7 kbp (Cell) trace remains non-stationary, with *α* = 1.17, consistent with anomalous subdiffusion in crowded media [58].

**Fig. 3.**
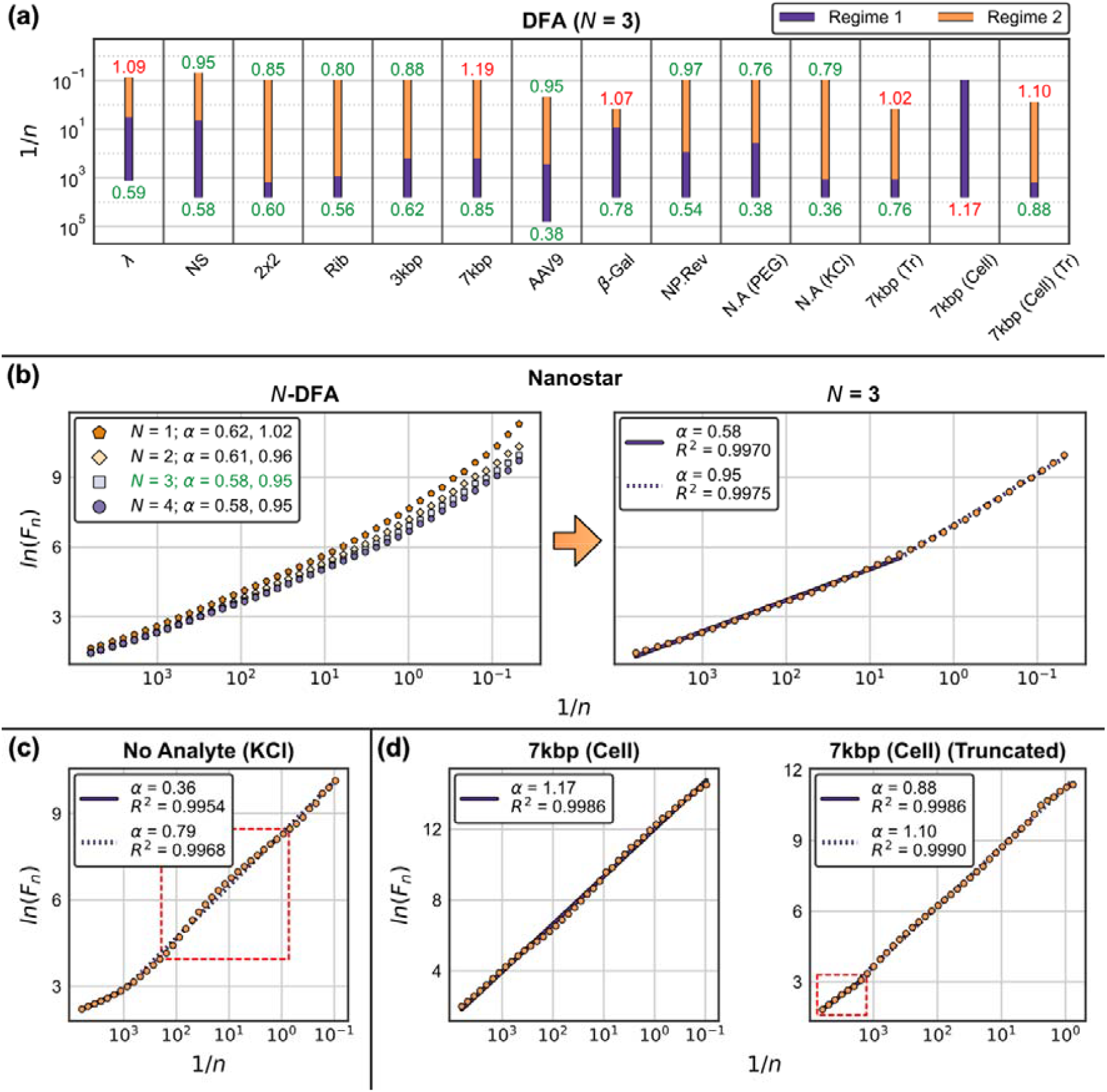
(a) DFA results for each trace, with values for each regime and red and green labels representing non-stationary and stationary scaling; (b) example self-similarity and dependence results for the Nanostar trace, illustrating how polynomial detrending orders were established (left plot, green label), and the resulting fitted regimes (right plot, purple lines); (c) large-scale regime (dotted purple line) nonlinearity in the No Analyte (KCl) trace, where the red square highlights the divergent scales; and (d) the emergence of the small-scale regime in the 7 kbp (Cell) trace following truncation (right plot, red square), demonstrating a behaviour consistent with the remaining traces. Abbreviations follow Fig. 2.

Truncation of the 7 kbp and 7 kbp (Cell) traces substantially improves scaling fits and reduces the degree of non-stationarity. In the 7 kbp trace, the large-scale exponent decreases from a 1.19 to a 1.02, and the small-scale exponent decreases from *α* = 0.85 to *α* = 0.76, with a higher crossover frequency and improved fit to the large-scale regime. Post-truncation, the scaling closely resembles that of the Ribosome and 2×2 traces but with higher exponents. Similarly, truncation of the 7 kbp (Cell) trace reduces a from 1.17 to 1.10, introduces a small-scale regime at *α* = 0.88, and improves the overall fit quality (Fig. 3(d)). These findings support the hypothesis that structural breaks are not inherent stochastic features and instead arise from external or deterministic influences, and when removed, reveal underlying scaling properties consistent with the remaining traces.

PSD analysis confirms that small-scale exponent reductions arise from an additional high-frequency regime, which lies beyond the resolution of DFA (Fig. S11). Although attenuated by the anti-aliasing filter, these regimes reflect underlying anti-persistent scaling and maintain scale-free properties up to the attenuation limit. Together, DFA and PSD provide complementary perspectives on variance scaling. While the PSD is sensitive to low-frequency trends and finite-size effects which can obscure large-scale exponents and crossovers in non-stationary contexts, it offers superior resolution of high-frequency (small-scale) structure beyond the numerical limits of DFA. By contrast, DFA suppresses polynomial trends by construction and is therefore more robust for estimating large-scale scaling in the presence of non-stationarities, the removal of segments, and remaining events [59], [60].

Overall, large-scale regime exponents span 0.76 <*α* < 1.19, while small-scale regime exponents range from 0.36 <*α* < 0.88, with most around *α* ≈0.6 (Fig. 3(a)). An additional third regime, observed in several traces, falls between. Notably, eight out of twelve traces exhibit stationary scaling across all regimes (, and all but the 7 kbp and 7 kbp (Cell) transition to stationary scaling before 10 Hz.

Examination of the stationarity and ensemble dependence properties demonstrates that every trace rejects the ADF null hypothesis of unit root across all segments, except for the 7 kbp and 7 kbp (Cell) traces (Fig. S13). The 7 kbp trace shows intermittent ADF assertion around two major structural breaks, while the 7 kbp (Cell) trace asserts more frequently as structural breaks progressively worsen. In contrast, the KPSS test results show greater variability within segment distributions (see Fig. S15), and trend stationary is rejected for the majority of segments. The only exception is the No Analyte (PEG) trace, which has the lowest regime exponents among the test set (Fig. 4(b)). Collectively, the combined ADF and KPSS results indicate that all traces exhibit fractionally integrated behaviour, with 0< *d* < 1, corresponding to 0.5 < *α* < 1.5.

**Fig. 4.**
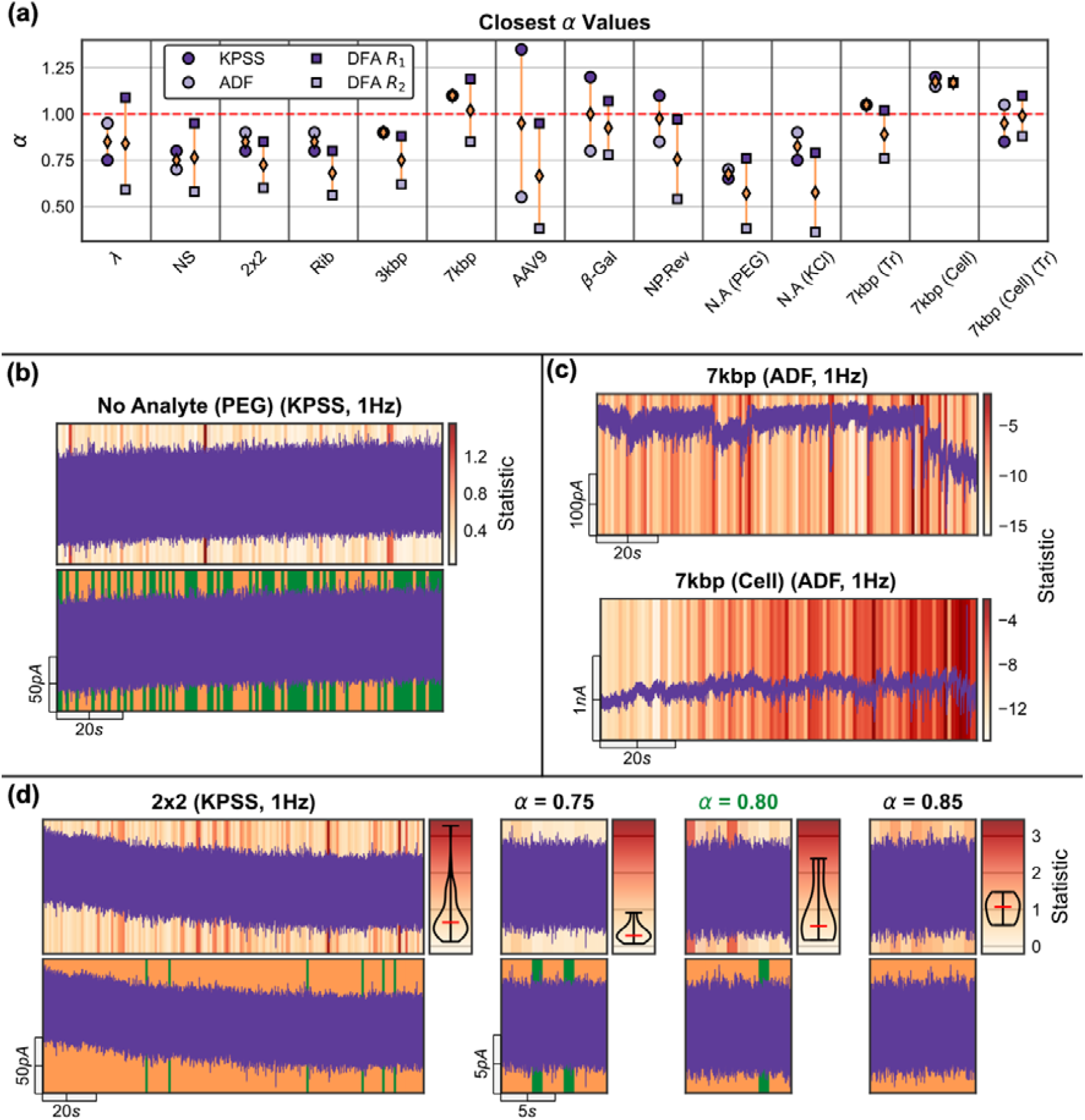
(a) Nearest reference set () values for each trace based on the KPSS and ADF statistic distributions, compared with the scaling exponents determined via DFA. The red line separates non-stationary from stationary scaling; (b) evolution of the KPSS statistic for the No Analyte (PEG) trace (top plot) and the corresponding test results (bottom plot); trend stationary assertion and rejection are indicated by green and orange overlays, respectively; (c) ADF statistic evolution for the 7 kbp (Cell) trace compared with the 7 kbp trace, which shows no evolution; and (d) a typical KPSS result alongside the three nearest reference set series based on the distribution of test statistics (adjacent violin plots), where the green label denotes the closest reference. Abbreviations follow Fig. 2.

Comparisons with the reference set, using EMD minimisation between the distributions of segment statistics and those of the reference set, shows that the mean of the closest fractional time-series exponents identified independently for KPSS and ADF correspond to the DFA scaling exponents (Fig. 4(a)). Specifically, the mean exponent either closely matches the average DFA exponent or reflects the large-scale DFA exponent, with a stronger bias toward the large-scale exponent as the regime crossover frequency increases. Although, due to limited segment numbers, both the AAV9 and β-galactosidase traces display larger discrepancies between their independently determined KPSS and ADF exponents. Nonetheless, the β-galactosidase remains close to the expected value, and the AAV9 correctly resolves the large-scale exponent. Moreover, the test statistic distributions for each trace are similar in shape to those from the reference set, further supporting consistency with fractional noise models (Fig. S12).

Truncation of the 7 kbp and 7 kbp (Cell) traces also makes them more stationary, consistent with the corrections observed in DFA. Specifically, while the non-truncated 7 kbp trace exhibits a regime crossover comparable to the 3 kbp trace, its mean exponent does not capture the large-scale regime. Truncation resolves this discrepancy, yielding an exponent that accurately reflects the large-scale behaviour. These results suggest that the original large-scale exponents in the non-truncated traces are inflated by structural breaks and that truncation reveals a more accurate representation of the underlying scaling behaviours.

Importantly, aside from the 7 kbp and 7 kbp (Cell), the temporal location of segments within each trace does not significantly influence the results (see Figs. S13-S20). No consistent patterns of assertion or rejection are observed, and segment statistics exhibit low variability. In the 7 kbp trace, variability in test outcomes is closely tied to the occurrence of structural breaks and where they occur within the segment. In contrast, the 7 kbp (Cell) trace shows a clear progression that is not solely linked to structural breaks (Fig. 4(c)). Overall, for the remaining traces, the temporal variability of the test statistics is comparable to that of the reference set, and no systematic evolution in stationarity or scaling is apparent.

In summary, second-order self-similarity and fractional scaling are consistently observed across all traces, with between one and three distinct scaling regimes ranging from anti-persistent to non-stationary. Notably, ten out of twelve traces exhibit stationary scaling across all regimes or at least above 10 Hz, with only one case of systematic deviation from self-similarity detected. Truncation of the 7 kbp and 7 kbp (Cell) traces reduces scaling exponents, improves fits, and enhances stationarity outcomes. Importantly, this reinforces the conclusion that structural breaks are quasi-deterministic artefacts, and truncation effectively isolates the underlying stochastic properties.

The combined KPSS and ADF analyses validate the presence of fractional integration across all traces and consistently resolve exponents, as confirmed by comparison with a reference set of fractional time-series. These exponents closely align with those derived from DFA and reflect their relative dominance across scaling regimes. Furthermore, KPSS and ADF outcomes demonstrate no significant temporal evolution in stationarity or dependence properties for any traces except those with structural breaks, indicating that these properties remain globally stable over time. However, the 7 kbp (Cell) trace exhibits additional non-stationary mechanisms that appear to underpin variance scaling alongside structural breaks. Finally, the distribution and temporal variability of segment-wise statistics closely resemble those of the reference set, providing significant qualitative evidence.

Overall, these findings support modelling within a fGn and fBm [57] framework, reflecting a superposition of fractional processes shaped by inherent system properties and experimental conditions.

In broader terms, this analysis demonstrates that deterministic trend stationarity is maintained across traces in the absence of structural breaks and non-stationary large-scale regimes, resulting in effectively stationary environments once low-frequency fluctuations are removed. For traces exhibiting large-scale non-stationarity, we show that most reach stationary scaling by 10 Hz. While these statistical properties are not strictly time-invariant, their temporal stability is sufficient to justify the assumption of covariance stationary for analytical purposes. We find that nonlinear detrending at an equivalent of 1 Hz is sufficient for the traces in our dataset, although lower cutoff frequencies may be suitable depending on the system characteristics. Nonetheless, maximising the detrending frequency is optimal, providing events are not attenuated. This minimises distortion of the relative offset due to correlation structure below the event scale, reduces the risk associated with non-stationary large-scale regimes, and improves the signal-to-noise ratio in a manner analogous to low-pass filtering.

Baseline trends are found to lie far outside the timescales associated with events in our testing set, and this separation is likely to generalise. However, overlap between the baseline and event scales remains a risk in nanopore sequencing applications, where event durations may span an order of magnitude longer. This further highlights the challenge posed by asymptotic tails, and how they can hinder accurate baseline determination.

Overall, our findings confirm that baseline fluctuations primarily reflect deterministic rather than stochastic features, consistent with prior visual inspection. Crucially, the effective removal of these trends is necessary to preserve the underlying stochastic properties that support valid statistical interpretation. This is consistent with the majority of nanopore event extraction algorithms, which prioritise baseline correction prior to thresholding [22], [23], [24], [25], [26], [61]. Our analysis supports this methodology and reiterates the importance of such removal. Regardless of the specific analysis pipeline, an effective removal of this component is required to regress the signal into a beneficial environment. Still, the field would benefit from the development of a predictor for baseline influence, which becomes particularly difficult under non-stationary large-scale regimes.

Despite these analyses demonstrating similar noise properties across the testing set, we identify asymptotic event tails specific to PEG traces, characterised as a scale-free monotonic decline to the baseline current. This complicates event endpoint identification as the gradual decline becomes entangled with steady-state noise, introducing potential biases in downstream analysis, and risking overlap with non-stationary baseline fluctuations. Even in strictly stationary environments, the issue persists as the canonical first baseline cross endpoint criterion becomes inherently governed by the noise variance, irrespective of the tail behaviour beyond the statistically expected endpoint.

While full-width-half-maximum has emerged as a common predictor of event duration, our analysis supports its use in PEG environments specifically due to endpoint ambiguity. However, this approach is not without bias, particularly in cases of slow and gradual decline. By comparison, the second-order differential method [21] is rendered ineffective, as the smooth and asymptotic nature, along with baseline overlap, fail to yield clear turning points. Conversely, KCl conditions are absent of asymptotic tails and remain compatible with both the canonical baseline cross criterion and the second-order differential method. This contrast highlights the need for either a dedicated event extraction algorithm tailored to PEG traces, or modifications to existing algorithms to account for this effect.

These challenges reflect broader properties of PEG signals. While the event amplification properties remain debated [1, 37], our findings raise important questions about the time required for the system to reach stability, and whether such effects should be explicitly considered during experimental design.

Taken together, several targeted refinements are warranted in nanopore analysis workflows. Scale-free, asymptotic event tails in PEG environments necessitate either dedicated extraction algorithms or modifications to existing methods to ensure accurate endpoint determination. While full-width-half-maximum remains a viable duration metric in such cases, its application must account for biases introduced by gradual tail decay. Likewise, effective baseline trend removal is critical to preserving the stochastic properties necessary for valid statistical interpretation, maintaining adherence to the proposed noise model, and ensuring assumptions in downstream analyses are met. Although maximising the detrending frequency is generally advantageous, the presence of asymptotic tails increases the risk of attenuating event components alongside the baseline. A preferable approach is to explicitly separate the true baseline from the signal without reliance on an estimated cutoff frequency, followed by correction of endpoint biases within the pure noise environment. Similarly, structural breaks must be detected and corrected to prevent spurious deviations in statistical descriptors and ensure consistency with the proposed model. Given the availability of effective change-point algorithms [27], such corrections can be incorporated into existing workflows with minimal modification, enabling statistically consistent handling of challenging traces. In practice, structural break correction should precede baseline determination, ensuring each source of bias is addressed independently. Collectively, these refinements define a clear pathway for reliable analysis of challenging traces.

Assessing multifractality is critical for determining whether the noise profiles and their respective scaling regimes exhibit monofractal (single exponent) behaviour or contain scaling heterogeneities arising from nonlinear effects. While prior distribution and dependence analyses establish fractional noise characteristics consistent with fGn and fBm, multifractality analysis examines whether fluctuations of different magnitudes exhibit distinct scaling laws, using higher-order moments (*q*-orders).

Multifractality may arise from nonlinear interactions that induce fluctuation-dependent correlations, manifesting as distinct scaling behaviours across fluctuation magnitudes. Such nonlinear mechanisms reflect complex dependence structures not captured by monofractal analysis. Additionally, this analysis differentiates intrinsic multifractality from artefacts such as structural breaks, variance shifts, or mixed noise sources, which produce spurious multifractal signatures. Finally, by examining changes in multifractal signatures following random shuffling, both scaling and multifractality can be attributed to temporal correlations or distributional effects (see Sec. II K 1 for a full breakdown). Overall, this analysis completes the characterisation of the stochastic and scaling properties. These findings corroborate prior distributional and dependence analyses, probe potential nonlinearities, and offer robust support for modelling the system within a fGn and fBm framework.

To characterise multifractality and its origins, MF-DFA [55] was applied to all events-removed traces and truncated versions of the 7 kbp and 7 kbp (Cell) traces. To assess whether multifractality arises from temporal correlations or distributional effects, MF-DFA was also performed on shuffled versions of each trace (see Sec. II K, Figs. S21-S22). Additionally T-DFA [56] was used to examine the temporal evolution of local scaling exponents *α*_*t*_ for pertinent scales in traces exhibiting multifractality (see Sec. II L, Figs. S23-S24). Together, these methods provided a comprehensive characterisation of multifractality, its origins, and its temporal evolution across all traces.

Analysis of MF-DFA results on the shuffled traces confirms that both scaling and multifractality are driven by temporal correlations rather than distributional properties, corroborating prior distributional findings. After shuffling, all *q*-order separation collapses and the scaling exponents *α*_*q*_ become consistent with uncorrelated white noise, demonstrating that heavy-tailed distributions do not contribute to these effects (Fig. 5(a)). Notably, in traces where structural breaks distort scaling and introduce multifractal signatures, these effects vanish upon shuffling. This reinforces that structural breaks are not impacting the noise distribution.

**Fig. 5.**
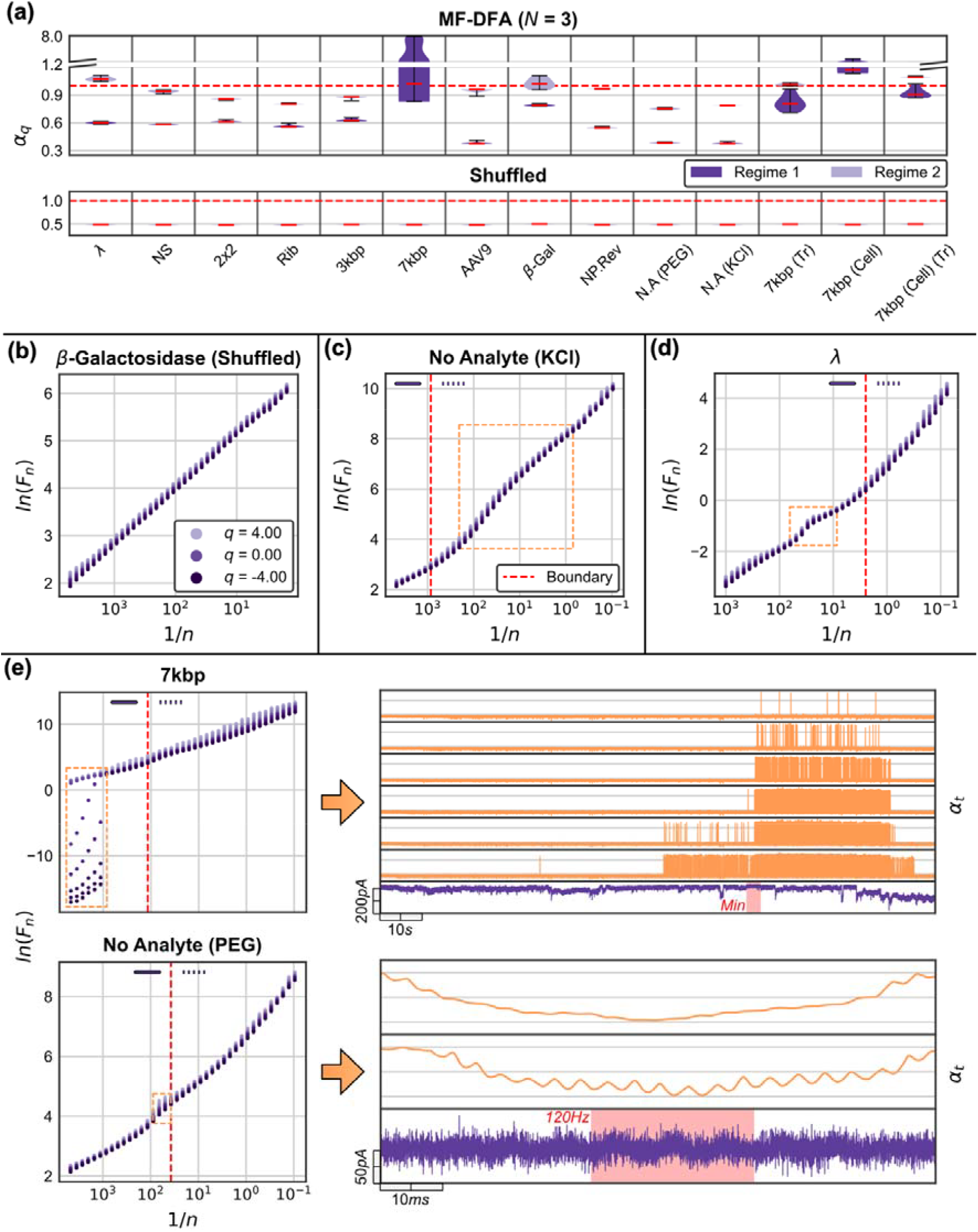
(a) Ensemble MF-DFA and shuffled MF-DFA results. The dotted red lines separate non-stationary from stationary scaling; (b) typical shuffled MF-DFA result showing uncorrelated scaling. Each of the eleven-orders is illustrated with sequential colouring; (c) -order alignment for the large-scale regime nonlinearity in the No Analyte (KCl) trace (orange box). The red line separates scaling regimes determined via DFA, labelled by their corresponding line style; (d) -order alignment for the trace line-frequency divergence (orange box); and (e) small-scale negative -order divergence in the 7 kbp trace and -order separation around line frequency for the No Analyte (PEG) trace both with the corresponding T-DFA of the divergent scales. The scales indicated by the orange boxes are displayed above each time-series, ordered from large (top) to small (bottom). For the 7 kbp trace, each scale spans **0.8** < ***α_t_*** < 2.2, and the red box highlights the minimum conductance point of the trace. For the No Analyte (PEG) trace, scales span **0.07** < ***α_t_*** < 0.33, and the red box highlights clear periodicity at 120 Hz. Abbreviations follow Fig. 2.

Considering the unshuffled results, several traces exhibit consistent monofractal behavior, characterised by parallel *q*-orders, constant *α*_*q*_, linear *α*_*q*_, and narrow multifractal spectra *D*_*q*_. These properties indicate that individual scaling regimes are well described by single-exponent, correlation-driven processes. Notably, the β-Galactosidase, 2×2 DNA origami, 30 nm AgNP (Reverse), No Analyte (KCl) and Nanostar traces display minimal deviation from this behaviour (Fig. S21).

Other traces remain monofractal but display localised effects. As previously established, both the *λ* and No Analyte (PEG) traces exhibit scale-specific line-frequency artifacts. In the *λ* trace, this appears as an additive, *q*-independent distortion that equally affects all fluctuation magnitudes (Fig. 5(d)). In contrast, the No Analyte (PEG) trace shows a *q*-dependent separation, suggesting interaction between intrinsic noise and periodic interference. T-DFA confirms intermittent 120 Hz modulation (Fig. 5(e)). Since these effects are confined in scale and fluctuation magnitude, and vanish upon shuffling, both traces remain globally monofractal. The AAV9 and 3 kbp traces similarly display localised positive *q*-order divergence without affecting overall monofractality. The AAV9 trace exhibits an abrupt divergence around 200 Hz that monotonically decreases, confirmed by T-DFA to arise from intermittent, unremoved events below the detection limit. In the 3 kbp trace, *q*-order separation occurs between 1-100 Hz and peaks near 10 Hz. T-DFA confirms that this effect arises from asymptotic PEG tails, with the peak separation corresponding with the largest asymptotic tail (Figs. S23-S24). In both cases, T-DFA reveals that increasing segment sizes progressively average out these contributions until their influence vanishes. The onset of separation aligns with the specific scales where these effects occur, and in the 3 kbp trace, a gradual increase reflects variability in the asymptotic tail lengths. The Ribosome trace exhibits consistent *q*-order separation in the large-scale regime, likely reflecting two competing fractional noise sources with similar scaling exponents, as supported by the collapse of separation upon shuffling. Additionally, extreme divergence at the largest scales is attributable to the singular structural break. Interestingly, the prominent bump observed in the No Analyte (KCl) trace is aligned across all *q*-orders, affecting small and large fluctuations equally without inducing divergence. Its removal upon shuffling suggests a correlation-driven feature, likely arising from system-intrinsic processes that do not introduce multifractal behaviour.

The 7 kbp and 7 kbp (Cell) traces exhibit extreme multifractality characterised by divergent negative *q*-orders at small scales and variable positive *q*-order separation across scales. These signatures arise from two interacting and intermittent mechanisms as confirmed by T-DFA (Fig. 5(e) and S23). First, structural breaks impact large fluctuations (*q* > 0), producing low local Hurst exponents *α*_*t*_. This effect non-uniformly impacts all scales due to the variable reversion rates. Second, smooth and highly correlated regions, reflecting intermittent dominance of the large-scale regime at the large/small-scale crossover, drive the divergence at negative *q*-orders through near-perfect local polynomial fits. In the 7 kbp trace, the onset of *q* < 0 divergence coincides with the minimum conductance state, whereas in the 7 kbp (Cell) trace it appears earlier during the conductance decline. This divergence resolves as the conductance recovers, first disappearing at larger scales before vanishing at smaller scales. Following the onset of multifractality, structural breaks intensify, resistive event sizes increase and translocation frequency rises, consistent with our prior visual analysis. Truncation effectively eliminates multifractality in both traces, eradicating negative *q*-order divergence and aligning positive *q*-order separation. T-DFA directly links the *q* > 0 effect to structural breaks, confirming that these fluctuations arise from transient system dynamics rather than intrinsic multifractality.

In conclusion, this analysis demonstrates that ten out of twelve traces exhibit primarily monofractal behavior, with only minor artefacts such as periodic interference, unremoved events or asymptotic tails. These results indicate that each dominant scaling regime is well described by a single effective exponent, even where multiple stochastic contributions may be present across different regimes. The remaining two traces, the 7 kbp and 7 kbp (Cell), display multifractality confined to small scales. This effect is directly linked to structural breaks, conductance states and highly correlated fluctuation states. Truncation eliminates these effects, restoring monofractality, reinforcing that multifractality is not an intrinsic stochastic feature but instead emerges from transient system dynamics. Importantly, these results corroborate the distribution and dependence analyses, reinforcing that nanopore noise is governed by temporal correlations rather than heavy-tailed distributions. Moreover, the combined evidence from distribution, self-similarity, dependence, and fractality analyses confirms that fGn and fBm provide appropriate models for characterising nanopore noise.

Through cross-validated methodology, Gaussianity emerges as a critical determinant in this study. Gaussian distributions underpin all analysed noise profiles, validating the choice of Gaussian-based statistical method and confirming that heavy-tailed alternatives, such as alpha-stable processes, are unsuitable for modelling these signals. Minor deviations, primarily asymptotic tails in event-associated directions and structural breaks, are attributable to event removal artifacts and quasi-deterministic first-order displacements, rather than intrinsic distributional properties. Direct confirmation of these divergences, including the preservation of higher-order moments, reinforce the assertion that Gaussian processes are optimal descriptors of nanopore noise.

Critically, multifractal analysis reveals that nanopore noise is driven primarily by temporal correlations rather than distributional properties, further ruling out extreme-value statistics and corroborating our previous findings. Apparent multifractal behaviour arises from transient system dynamics and, in the case of structural breaks, disappears upon truncation. Targeted T-DFA analysis confirms the exact deterministic origin of these signatures, attributable to line frequency interference, event removal artefacts, structural breaks, and associated fluctuation states. Thus, dependence structures identified via second-order analyses, including R/S, DFA and PSD, reveal multiple monofractal scaling regimes corresponding to distinct fractional processes, ranging from 0< *α* < 1.19. Small-scale regimes likely result from exponents below the conventional range (*α* < 0), shaped by the effects of anti-aliasing filters, but nonetheless exhibit behaviour consistent with regimes within the conventional range. Moreover, our novel KPSS/ADF based analysis, utilising comparisons with a reference set of fractional time-series, confirm that correlation and time evolution properties directly align with fractional noise models. Although the 7 kbp (Cell) trace exhibits evolving dependence effects, this reflects an evolving non-stationary scaling exponent, opposed to a distinct process to the remainder of the set. Modelling regime evolution lies beyond the scope of this manuscript, and further work would be necessary to fully capture this noise profile.

Multiple analyses show that structural breaks do not represent intrinsic stochastic features but instead arise from deterministic analyte interactions. This interpretation is strongly supported by the observed links between reversible conductance states, conductance trends, preservation of higher-order distributional properties, translocating analyte signatures, correlative excitations, and multifractal behaviour. Further evidence from truncated traces shows that divergent behaviour associated with structural breaks reverts upon removal, restoring statistical properties aligned with the remaining traces. This confirms that such features are non-intrinsic and primarily reflect first-order displacement effects, rather than modifications to the underlying stochastic process.

This behaviour mirrors stepwise nanopore event signatures, where algorithms such as CUSUM [28], [29], [62] operate on this mean displacement criteria within Gaussian noise. We demonstrate that such displacements can occur in the absence of translocation events and hypothesise that they originate from analyte activity in the access region, such as absorption, blockage, or aggregation. These effects are consistent with modulation of the pore resistance and local charge density due to analyte aggregation, given that primary sources of long-range dependent noise originate in the access region [14].

Analyses such as KPSS/ADF reference comparisons, DFA, MF-DFA, and T-DFA remain rarely applied in the nanopore field, yet they yield rich, directly interpretable insights into noise behaviour. Demonstrating their utility here should encourage their broader adoption, enabling more accurate and statistically grounded characterisation of nanopore signals. This is particularly critical for analyte-containing recordings, where noise stability is reduced relative to analyte-free conditions and rigorous methodology is required to detect and account for potential divergences, structural breaks, and other non-idealities. For a practical step-by-step guide to conducting the analyses presented in this manuscript, see Supplemental Material under the heading *Practical Analysis Pipeline for Trace Validation*.

## IV. CONCLUSIONS

This study demonstrated that nanopore noise can be modelled as a superposition of multiple fGn and fBm, contingent upon the assumption of underlying Gaussian distributions. Using a broad set of independently acquired nanopore traces, we demonstrate consistent behaviour across varied experimental conditions that validates this assumption. We also identify significant implications arising from baseline influences, asymptotic event tails, and structural breaks. Importantly, we investigate and classify individual deviations from the required statistical properties, establishing a rigorous empirical foundation for the model.

While the multi-regime model is well-known to mechanistic nanopore noise studies [12], [13], [14], [15], prior work has not established the conditions necessary to enforce a composite fractional model, nor linked second-order exponents to interpretable model parameters as demonstrated here. This formal validation enables precise and meaningful interpretation of these exponents, facilitating comparison across independent experiments and systems, while disentangling intrinsic noise properties from transient and artefactual features. In practical terms, it allows algorithms, formulated in statistical language, to directly target the statistical descriptors defined by the validated model. Although numerous approaches exist for filtering, event extraction, and higher-level signal processing, many lack a precise definition of the statistical properties they aim to preserve or estimate and often rely on methods unsuited to the underlying noise characteristics. By confirming the Gaussian nature of nanopore noise and linking second-order scaling exponents to interpretable parameters within a composite fractional model, this work enables the principled design of algorithms that address well-defined statistical objectives.

Despite event specific differences, no fundamental distinctions are observed in the analyte-free and analyte-containing environments. Although the analyte-free KCl condition exhibits a systematic deviation from self-similarity in the large-scale regime, this feature has also been reported in PEG traces and attributed to absorption effects [30], suggesting a shared underlying mechanism. Furthermore, no major distributional or dependence differences distinguish analyte-free and analyte-containing traces, reinforcing that analyte presence does not fundamentally alter nanopore noise beyond regime exponents. This is consistent with the expected influence of increased charge within the system, given the known origin of major noise sources [14].

Traces containing structural breaks are typically disregarded and re-recorded under modified experimental conditions. This practice is common in both analyte signature-based inference and mechanistic noise studies. In this work, we deliberately include such challenging traces to examine the implications of structural breaks. We identify these breaks as quasi-deterministic phenomena, characterised by abrupt mean shifts and subsequent reversions. A single non-reverting case is observed in one trace, but its permanent nature and lack of effect on other statistical properties render it of limited further relevance. These findings have important implications as the field moves towards the analysis of more challenging analytes, where such phenomena may be unavoidable, or re-recording is infeasible. Provided that analytical pipelines account for the presence of these breaks, the underlying noise behaviour remains unchanged, suggesting minimal modification to existing analysis workflows is required.

Finally, these findings enable principled reconstruction and generation of realistic noise profiles using established fGn and fBm synthesis methods [51], [63], [64], [65], subject to efficient extraction of the model parameters under a nonlinear formulation. This capability not only supports the creation of high-fidelity performance evaluation datasets for signal processing, but also the controlled generation and augmentation of data for machine learning while maintaining statistical fidelity to experimental conditions.

## VI. ACKNOWLEDGEMENTS

The authors would like to thank Dr James Yates (**ITQB NOVA, Lisbon, PT**) for kindly sharing the λ-DNA dataset and Dr Michelangelo Paci and Dr Alessandro Porro (Elements srl) for sharing the AAV9 datasets. C.C.C.C., C.W., and P.A acknowledge funding from the Engineering and Physical Sciences Research Council UK (EPSRC) Healthcare Technologies for the grant EP/W004933/1, and C.C.C.C., P.A. acknowledge the Biotechnology and Biological Sciences Research Council (BBSRC) for the grant BB/X003086/1. D.C. acknowledges funding from the EPSRC EP/W524372/1 and the Bragg Centre for Materials Research.

## VII. DATA AVAILABILITY

Data supporting this work can be freely accessed via the University of Leeds repository: https://doi.org/10.5518/1795

## REFERENCES

[1] F. Marcuccio et al., ‘Mechanistic Study of the Conductance and Enhanced Single-Molecule Detection in a Polymer–Electrolyte Nanopore’, ACS Nanoscience Au, vol. 3, no. 2. pp. 172–181, 2023. doi: 10.1021/acsnanoscienceau.2c00050.

[2] C. Koch et al., ‘Nanopore sequencing of DNA-barcoded probes for highly multiplexed detection of microRNA, proteins and small biomarkers’, Nat. Nanotechnol., vol. 18, no. 12, pp. 1483–1491, Dec. 2023, doi: 10.1038/s41565-023-01479-z.

[3] C. C. Chau, S. E. Radford, E. W. Hewitt, and P. Actis, ‘Macromolecular Crowding Enhances the Detection of DNA and Proteins by a Solid-State Nanopore’, Nano Lett., vol. 20, no. 7, pp. 5553–5561, Jul. 2020, doi: 10.1021/acs.nanolett.0c02246.

[4] C. Chau et al., ‘Probing RNA Conformations Using a Polymer-Electrolyte Solid-State Nanopore’, ACS Nano, vol. 16, no. 12, pp. 20075–20085, Dec. 2022, doi: 10.1021/acsnano.2c08312.

[5] S. Confederat et al., ‘Next-Generation Nanopore Sensors Based on Conductive Pulse Sensing for Enhanced Detection of Nanoparticles’, Small, vol. 20, no. 4, p. 2305186, 2024, doi: 10.1002/smll.202305186.

[6] L. Xue, H. Yamazaki, R. Ren, M. Wanunu, A. P. Ivanov, and J. B. Edel, ‘Solid-state nanopore sensors’, Nat. Rev. Mater., vol. 5, no. 12, pp. 931–951, Dec. 2020, doi: 10.1038/s41578-020-0229-6.

[7] R. Ren et al., ‘Single-Molecule Binding Assay Using Nanopores and Dimeric NP Conjugates’, Adv. Mater., vol. 33, no. 38, p. 2103067, Sep. 2021, doi: 10.1002/adma.202103067.

[8] S. Cai et al., ‘Single-molecule amplification-free multiplexed detection of circulating microRNA cancer biomarkers from serum’, Nat. Commun., vol. 12, no. 1, p. 3515, Jun. 2021, doi: 10.1038/s41467-021-23497-y.

[9] F. Bošković and U. F. Keyser, ‘Nanopore microscope identifies RNA isoforms with structural colours’, Nat. Chem., vol. 14, no. 11, pp. 1258–1264, Nov. 2022, doi: 10.1038/s41557-022-01037-5.

[10] N. A. W. Bell and U. F. Keyser, ‘Digitally encoded DNA nanostructures for multiplexed, single-molecule protein sensing with nanopores’, Nat. Nanotechnol., vol. 11, no. 7, pp. 645–651, Jul. 2016, doi: 10.1038/nnano.2016.50.

[11] A. Dorey and S. Howorka, ‘Nanopore DNA sequencing technologies and their applications towards single-molecule proteomics’, Nat. Chem., vol. 16, no. 3, pp. 314–334, Mar. 2024, doi: 10.1038/s41557-023-01322-x.

[12] R. M. M. Smeets, U. F. Keyser, N. H. Dekker, and C. Dekker, ‘Noise in solid-state nanopores’, Proceedings of the National Academy of Sciences, vol. 105, no. 2. pp. 417–421, 2008. doi: 10.1073/pnas.0705349105.

[13] S. Liang et al., ‘Noise in nanopore sensors: Sources, models, reduction, and benchmarking’, Nanotechnology and Precision Engineering, vol. 3, no. 1. pp. 9–17, 2020. doi: 10.1016/j.npe.2019.12.008.

[14] C. Wen et al., ‘Generalized Noise Study of Solid-State Nanopores at Low Frequencies’, ACS Sensors, vol. 2, no. 2. pp. 300–307, 2017. doi: 10.1021/acssensors.6b00826.

[15] V. Tabard-Cossa, D. Trivedi, M. Wiggin, N. N. Jetha, and A. Marziali, ‘Noise analysis and reduction in solid-state nanopores’, Nanotechnology, vol. 18, no. 30, p. 305505, Jun. 2007, doi: 10.1088/0957-4484/18/30/305505.

[16] J. D. Uram, K. Ke, and M. Mayer, ‘Noise and Bandwidth of Current Recordings from Submicrometer Pores and Nanopores’, ACS Nano, vol. 2, no. 5, pp. 857–872, May 2008, doi: 10.1021/nn700322m.

[17] S. F. Knowles, U. F. Keyser, and A. L. Thorneywork, ‘Noise properties of rectifying and non-rectifying nanopores’, Nanotechnology, vol. 31, no. 10, p. 10LT01, Mar. 2020, doi: 10.1088/1361-6528/ab5be3.

[18] M. Zorkot, R. Golestanian, and D. J. Bonthuis, ‘The Power Spectrum of Ionic Nanopore Currents: The Role of Ion Correlations’, Nano Lett., vol. 16, no. 4, pp. 2205–2212, Apr. 2016, doi: 10.1021/acs.nanolett.5b04372.

[19] A. Fragasso, S. Schmid, and C. Dekker, ‘Comparing Current Noise in Biological and Solid-State Nanopores’, ACS Nano, vol. 14, no. 2, pp. 1338–1349, Feb. 2020, doi: 10.1021/acsnano.9b09353.

[20] R. Gao, M. A. Edwards, J. M. Harris, and H. S. White, ‘Shot noise sets the limit of quantification in electrochemical measurements’, Curr. Opin. Electrochem., vol. 22, pp. 170–177, Aug. 2020, doi: 10.1016/j.coelec.2020.05.010.

[21] Z. Gu, Y.-L. Ying, C. Cao, P. He, and Y.-T. Long, ‘Accurate Data Process for Nanopore Analysis’, Analytical Chemistry, vol. 87, no. 2. pp. 907–913, 2015. doi: 10.1021/ac5028758.

[22] Z. Sun et al., ‘AutoNanopore: An Automated Adaptive and Robust Method to Locate Translocation Events in Solid-State Nanopore Current Traces’, ACS Omega, vol. 7, no. 42. pp. 37103–37111, 2022. doi: 10.1021/acsomega.2c02927.

[23] C. Plesa and C. Dekker, ‘Data analysis methods for solid-state nanopores’, Nanotechnology, vol. 26, no. 8. p. 084003, 2015. doi: 10.1088/0957-4484/26/8/084003.

[24] H.-F. Wang et al., ‘Real-time Event Recognition and Analysis System for Nanopore Study’, Chin. J. Anal. Chem., vol. 46, no. 6, pp. 843–850, Jun. 2018, doi: 10.1016/S1872-2040(18)61090-4.

[25] D. Pedone, M. Firnkes, and U. Rant, ‘Data Analysis of Translocation Events in Nanopore Experiments’, Analytical Chemistry, vol. 81, no. 23. pp. 9689–9694, 2009. doi: 10.1021/ac901877z.

[26] Y. M. N. D. Y. Bandara, J. Saharia, B. I. Karawdeniya, P. Kluth, and M. J. Kim, ‘Nanopore Data Analysis: Baseline Construction and Abrupt Change-Based Multilevel Fitting’, Analytical Chemistry, vol. 93, no. 34. pp. 11710–11718, 2021. doi: 10.1021/acs.analchem.1c01646.

[27] C. Wen, D. Dematties, and S.-L. Zhang, ‘A Guide to Signal Processing Algorithms for Nanopore Sensors’, ACS Sensors, vol. 6, no. 10. pp. 3536–3555, 2021. doi: 10.1021/acssensors.1c01618.

[28] J. H. Forstater et al., ‘MOSAIC: A Modular Single-Molecule Analysis Interface for Decoding Multistate Nanopore Data’, Analytical Chemistry, vol. 88, no. 23. pp. 11900–11907, 2016. doi: 10.1021/acs.analchem.6b03725.

[29] C. Raillon, P. Granjon, M. Graf, L. J. Steinbock, and A. Radenovic, ‘Fast and automatic processing of multi-level events in nanopore translocation experiments’, Nanoscale, vol. 4, no. 16. p. 4916, 2012. doi: 10.1039/c2nr30951c.

[30] S. F. Knowles, N. E. Weckman, V. J. Y. Lim, D. J. Bonthuis, U. F. Keyser, and A. L. Thorneywork, ‘Current Fluctuations in Nanopores Reveal the Polymer-Wall Adsorption Potential’, Phys. Rev. Lett., vol. 127, no. 13, p. 137801, Sep. 2021, doi: 10.1103/PhysRevLett.127.137801.

[31] S. Gravelle, R. R. Netz, and L. Bocquet, ‘Adsorption Kinetics in Open Nanopores as a Source of Low-Frequency Noise’, Nano Lett., vol. 19, no. 10, pp. 7265–7272, Oct. 2019, doi: 10.1021/acs.nanolett.9b02858.

[32] D. Dematties, C. Wen, and S.-L. Zhang, ‘A Generalized Transformer-Based Pulse Detection Algorithm’, ACS Sensors, vol. 7, no. 9. pp. 2710–2720, 2022. doi: 10.1021/acssensors.2c01218.

[33] D. Dematties, C. Wen, M. D. Pérez, D. Zhou, and S.-L. Zhang, ‘Deep Learning of Nanopore Sensing Signals Using a Bi-Path Network’, ACS Nano, vol. 15, no. 9. pp. 14419–14429, 2021. doi: 10.1021/acsnano.1c03842.

[34] C. Wen, S. Zeng, Z. Zhang, K. Hjort, R. Scheicher, and S.-L. Zhang, ‘On nanopore DNA sequencing by signal and noise analysis of ionic current’, Nanotechnology, vol. 27, no. 21, p. 215502, Apr. 2016, doi: 10.1088/0957-4484/27/21/215502.

[35] R. A. Al-Waqfi, C. J. Khan, O. J. Irving, L. Matthews, and T. Albrecht, ‘Crowding Effects during DNA Translocation in Nanopipettes’, ACS Nano, vol. 19, no. 17, pp. 16803–16812, May 2025, doi: 10.1021/acsnano.5c01529.

[36] S. W. Kowalczyk, A. Y. Grosberg, Y. Rabin, and C. Dekker, ‘Modeling the conductance and DNA blockade of solid-state nanopores’, Nanotechnology, vol. 22, no. 31, p. 315101, Jul. 2011, doi: 10.1088/0957-4484/22/31/315101.

[37] E. G. Agyemang et al., ‘Multimodal nanoparticle analysis enabled by a polymer electrolyte nanopore combined with nanoimpact electrochemistry’, Faraday Discuss., vol. 257, no. 0, pp. 303–315, 2025, doi: 10.1039/D4FD00143E.

[38] C. C. Chau, C. M. Maffeo, A. Aksimentiev, S. E. Radford, E. W. Hewitt, and P. Actis, ‘Single molecule delivery into living cells’, Nat. Commun., vol. 15, no. 1, p. 4403, May 2024, doi: 10.1038/s41467-024-48608-3.

[39] S. Confederat, I. Sandei, G. Mohanan, C. Wälti, and P. Actis, ‘Nanopore fingerprinting of supramolecular DNA nanostructures’, Biophysical Journal, vol. 121, no. 24. pp. 4882–4891, 2022. doi: 10.1016/j.bpj.2022.08.020.

[40] J. P. Fried et al., ‘In situ solid-state nanopore fabrication’, Chem. Soc. Rev., vol. 50, no. 8, pp. 4974–4992, Apr. 2021, doi: 10.1039/D0CS00924E.

[41] J. P. Fried et al., ‘Localised solid-state nanopore fabrication via controlled breakdown using on-chip electrodes’, Nano Res., vol. 15, no. 11, pp. 9881–9889, Nov. 2022, doi: 10.1007/s12274-022-4535-8.

[42] S. S. Shapiro and M. B. Wilk, ‘An analysis of variance test for normality (complete samples)†’, Biometrika, vol. 52, no. 3–4, pp. 591–611, Dec. 1965, doi: 10.1093/biomet/52.3-4.591.

[43] H. E. Hurst, ‘Long-Term Storage Capacity of Reservoirs’, Trans. Am. Soc. Civ. Eng., vol. 116, no. 1, pp. 770–799, Jan. 1951, doi: 10.1061/TACEAT.0006518.

[44] B. B. Mandelbrot and J. R. Wallis, ‘Robustness of the rescaled range R/S in the measurement of noncyclic long run statistical dependence’, Water Resour. Res., vol. 5, no. 5, pp. 967–988, 1969, doi: 10.1029/WR005i005p00967.

[45] C.-K. Peng, S. V. Buldyrev, S. Havlin, M. Simons, H. E. Stanley, and A. L. Goldberger, ‘Mosaic organization of DNA nucleotides’, Phys. Rev. E, vol. 49, no. 2, pp. 1685–1689, Feb. 1994, doi: 10.1103/PhysRevE.49.1685.

[46] R. Weron, ‘Estimating long-range dependence: finite sample properties and confidence intervals’, Phys. Stat. Mech. Its Appl., vol. 312, no. 1, pp. 285–299, Sep. 2002, doi: 10.1016/S0378-4371(02)00961-5.

[47] S. Bianchi, ‘fathon: A Python package for a fast computation of detrendend fluctuation analysis and related algorithms’, J. Open Source Softw., vol. 5, no. 45, p. 1828, Jan. 2020, doi: 10.21105/joss.01828.

[48] J. W. Kantelhardt, E. Koscielny-Bunde, H. H. A. Rego, S. Havlin, and A. Bunde, ‘Detecting long-range correlations with detrended fluctuation analysis’, Phys. Stat. Mech. Its Appl., vol. 295, no. 3, pp. 441–454, Jun. 2001, doi: 10.1016/S0378-4371(01)00144-3.

[49] D. Kwiatkowski, P. C. B. Phillips, P. Schmidt, and Y. Shin, ‘Testing the null hypothesis of stationarity against the alternative of a unit root: How sure are we that economic time series have a unit root?’, J. Econom., vol. 54, no. 1, pp. 159–178, Oct. 1992, doi: 10.1016/0304-4076(92)90104-Y.

[50] D. A. Dickey and W. A. Fuller, ‘Likelihood Ratio Statistics for Autoregressive Time Series with a Unit Root’, Econometrica, vol. 49, no. 4, pp. 1057–1072, 1981, doi: 10.2307/1912517.

[51] R. B. Davies and D. S. Harte, ‘Tests for Hurst effect’, Biometrika, vol. 74, no. 1, pp. 95–101, Mar. 1987, doi: 10.1093/biomet/74.1.95.

[52] D. Lee and P. Schmidt, ‘On the power of the KPSS test of stationarity against fractionally-integrated alternatives’, J. Econom., vol. 73, no. 1, pp. 285–302, Jul. 1996, doi: 10.1016/0304-4076(95)01741-0.

[53] U. Hassler and J. Wolters, ‘On the power of unit root tests against fractional alternatives’, Econ. Lett., vol. 45, no. 1, pp. 1–5, May 1994, doi: 10.1016/0165-1765(94)90049-3.

[54] B. Hobijn, P. H. Franses, and M. Ooms, ‘Generalizations of the KPSS-test for stationarity’, Stat. Neerlandica, vol. 58, no. 4, pp. 483–502, 2004, doi: 10.1111/j.1467-9574.2004.00272.x.

[55] J. W. Kantelhardt, S. A. Zschiegner, E. Koscielny-Bunde, S. Havlin, A. Bunde, and H. E. Stanley, ‘Multifractal detrended fluctuation analysis of nonstationary time series’, Phys. Stat. Mech. Its Appl., vol. 316, no. 1, pp. 87–114, Dec. 2002, doi: 10.1016/S0378-4371(02)01383-3.

[56] E. A. Ihlen, ‘Introduction to Multifractal Detrended Fluctuation Analysis in Matlab’, Front. Physiol., vol. 3, Jun. 2012, doi: 10.3389/fphys.2012.00141.

[57] B. B. Mandelbrot and J. W. Van Ness, ‘Fractional brownian motions, fractional noises and applications’, SIAM Rev., vol. 10, no. 4, pp. 422–437, 1968, doi: 10.1137/1010093.

[58] M. Weiss, M. Elsner, F. Kartberg, and T. Nilsson, ‘Anomalous Subdiffusion Is a Measure for Cytoplasmic Crowding in Living Cells’, Biophys. J., vol. 87, no. 5, pp. 3518–3524, Nov. 2004, doi: 10.1529/biophysj.104.044263.

[59] Z. Chen, P. Ch. Ivanov, K. Hu, and H. E. Stanley, ‘Effect of nonstationarities on detrended fluctuation analysis’, Phys. Rev. E, vol. 65, no. 4, p. 041107, Apr. 2002, doi: 10.1103/PhysRevE.65.041107.

[60] N. W. Churchill et al., ‘The suppression of scale-free fMRI brain dynamics across three different sources of effort: aging, task novelty and task difficulty’, Sci. Rep., vol. 6, no. 1, p. 30895, Aug. 2016, doi: 10.1038/srep30895.

[61] J. Järlebark, W. Liu, A. Shaji, J. Sha, and A. Dahlin, ‘Solid-State Nanopore Sensors: Analyte Quantification by Event Frequency Analysis at High Voltages’, Anal. Chem., vol. 97, no. 8, pp. 4359–4364, Mar. 2025, doi: 10.1021/acs.analchem.4c05037.

[62] P. Granjon, ‘The CUSUM algorithm a small review’, GIPSA-lab, Report, Mar. 2014.

[63] A. T. A. Wood and G. Chan, ‘Simulation of Stationary Gaussian Processes in [0, 1] d’, J. Comput. Graph. Stat., vol. 3, no. 4, pp. 409–432, Dec. 1994, doi: 10.1080/10618600.1994.10474655.

[64] J. R. M. Hosking, ‘Modeling persistence in hydrological time series using fractional differencing’, Water Resour. Res., vol. 20, no. 12, pp. 1898–1908, 1984, doi: 10.1029/WR020i012p01898.

[65] D. B. Percival, ‘Exact simulation of complex-valued Gaussian stationary processes via circulant embedding’, Signal Process., vol. 86, no. 7, pp. 1470–1476, Jul. 2006, doi: 10.1016/j.sigpro.2005.08.003.

