## Supporting Information for "Empirical Validation of Composite Fractional Noise Models in Nanopore Signals"

### for

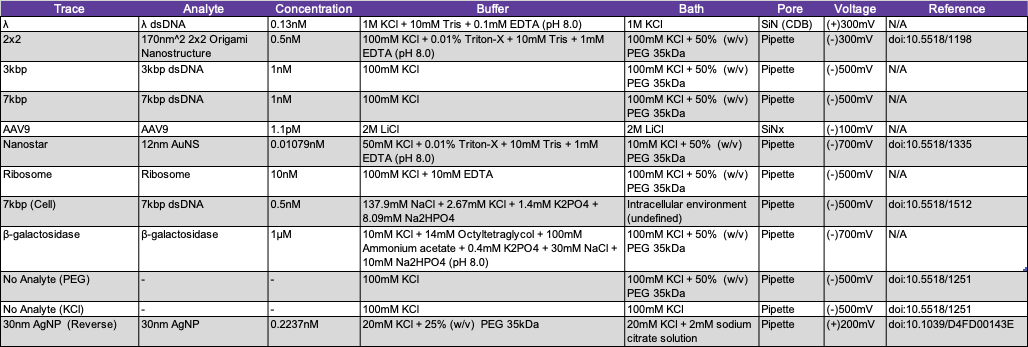

Table SI. Experimental details of the twelve traces used in this work. Further details are provided in the Methods section of the main text.

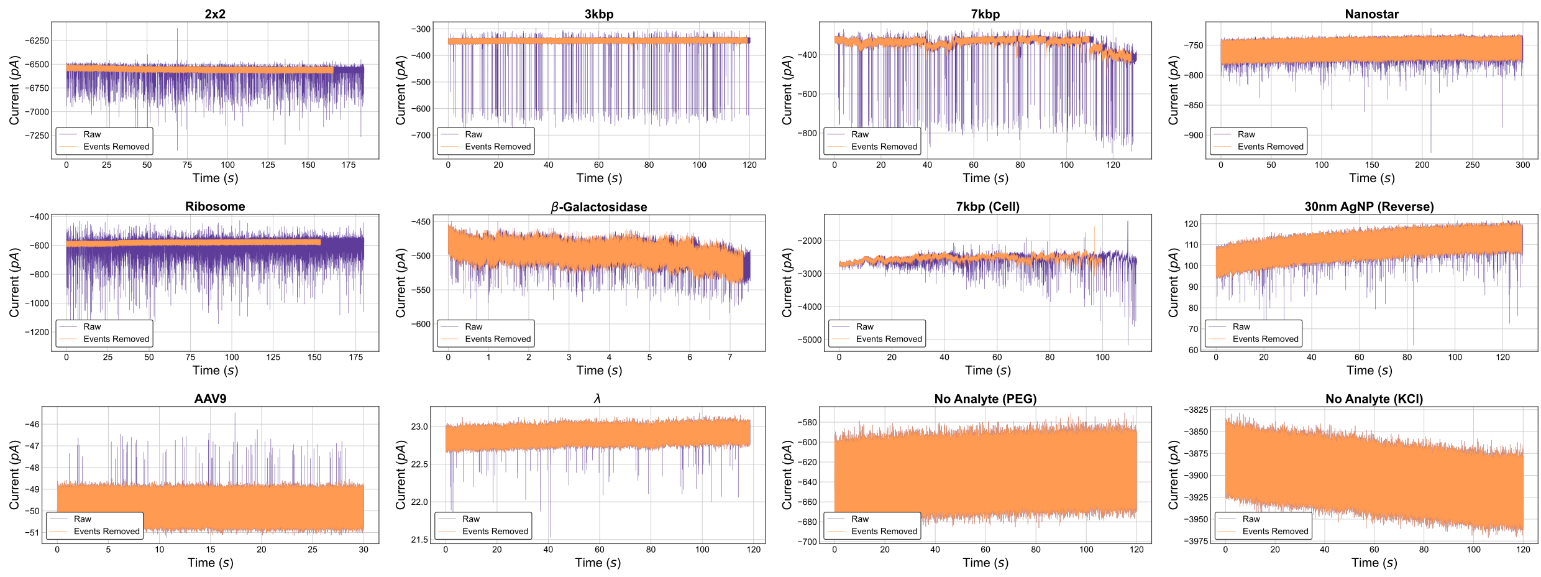
Fig. S1. The twelve traces before and after event removal as described in the Methods section of the main text.

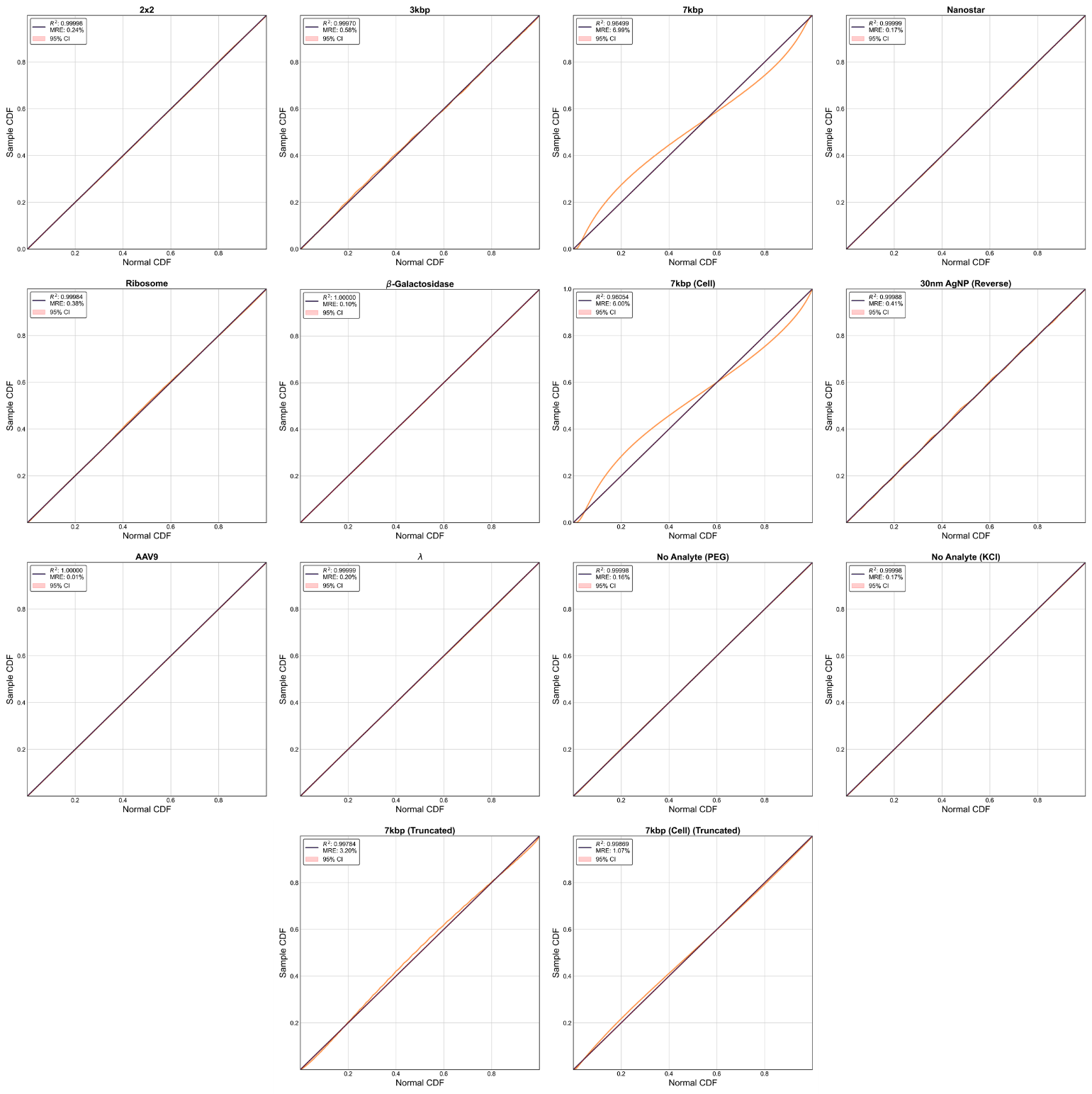
Fig. S2. PP plots comparing trace distributions to normal distributions. Methodology is provided in the Methods section of the main text.

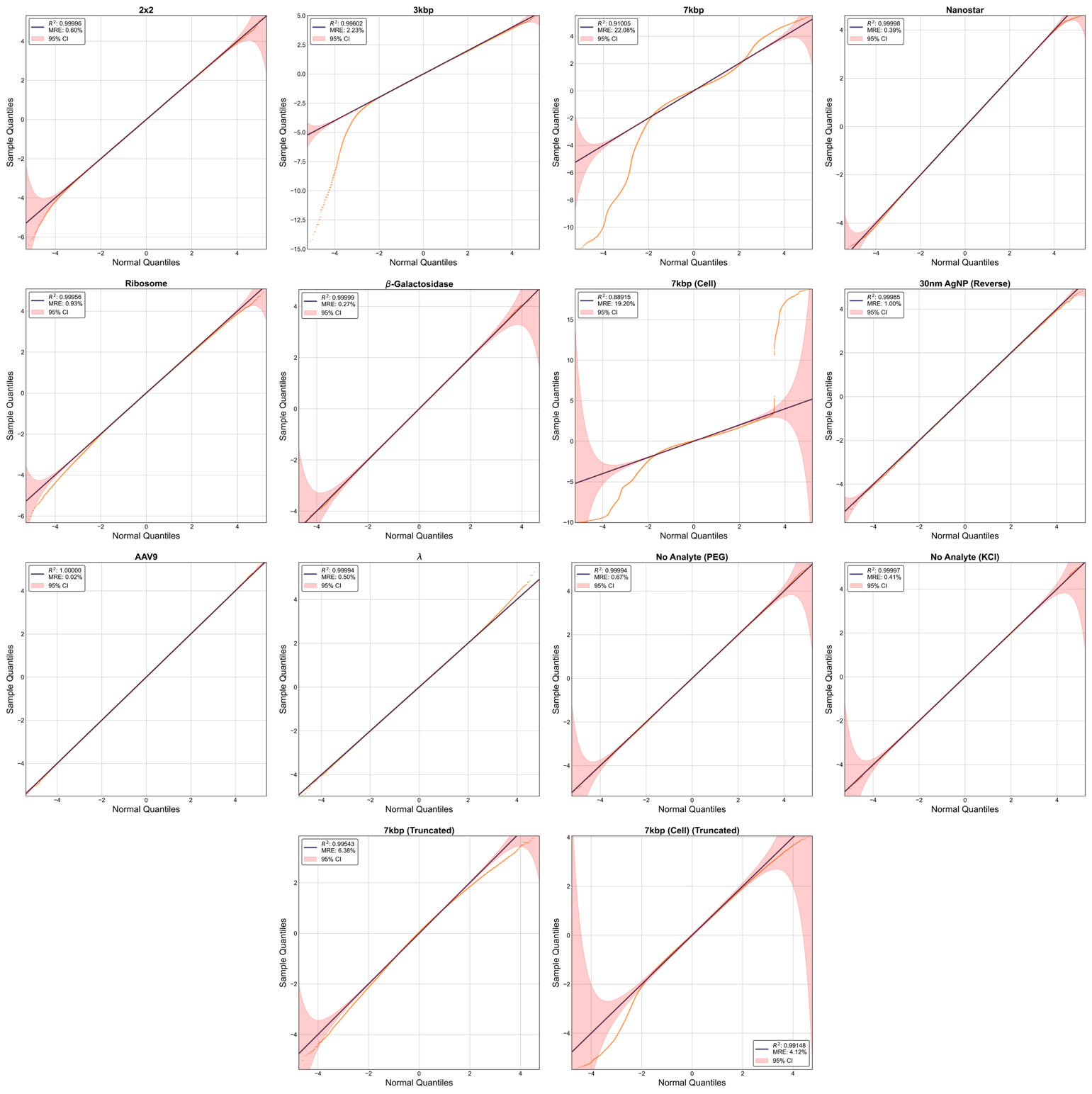
Fig. S3. QQ plots comparing trace distributions to normal distributions. Methodology is provided in the Methods section of the main text.

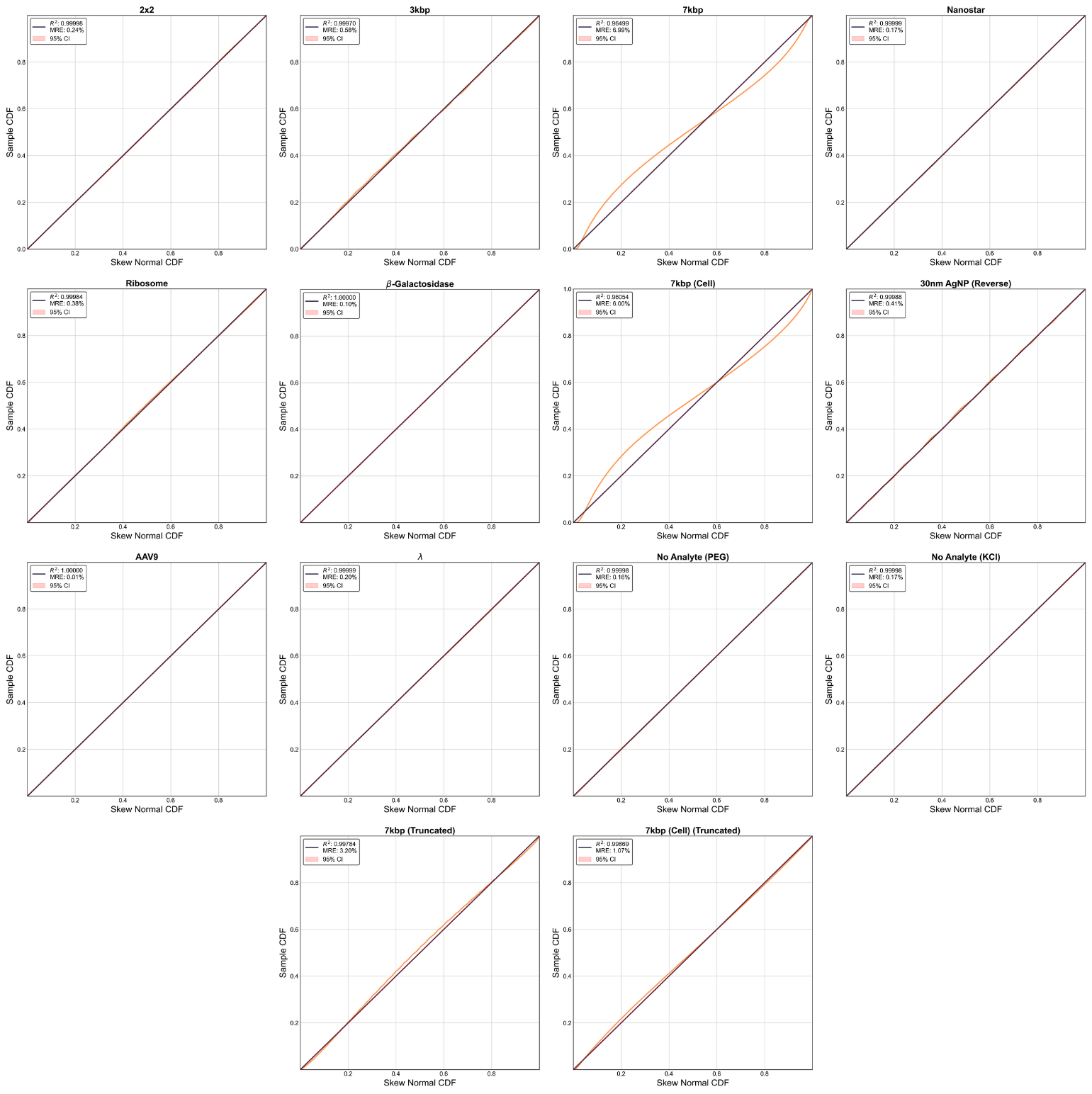
Fig. S4. PP plots comparing trace distributions to skew normal distributions. Methodology is provided in the Methods section of the main text.

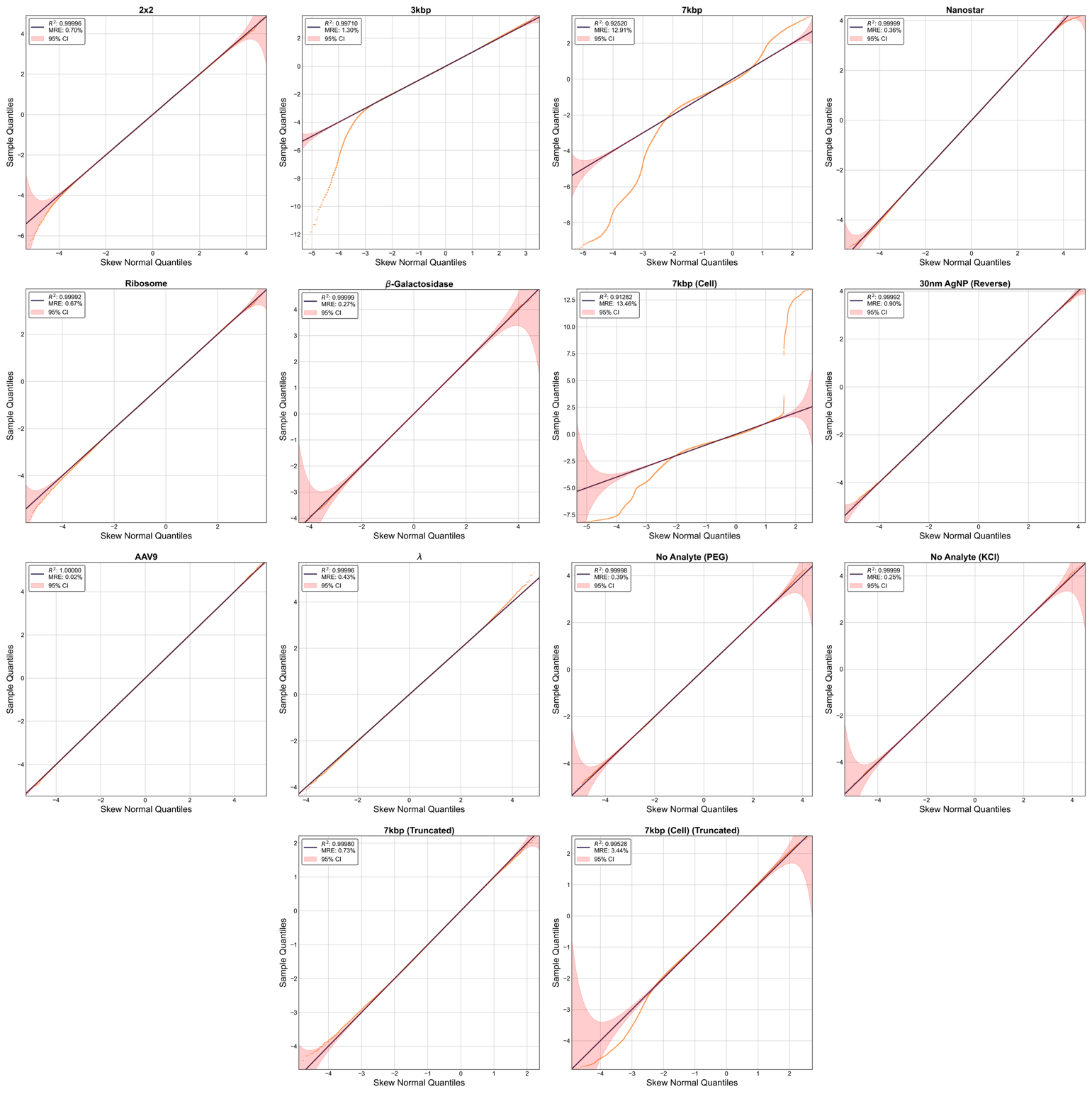
Fig. S5. QQ plots comparing trace distributions to skew normal distributions. Methodology is provided in the Methods section of the main text.

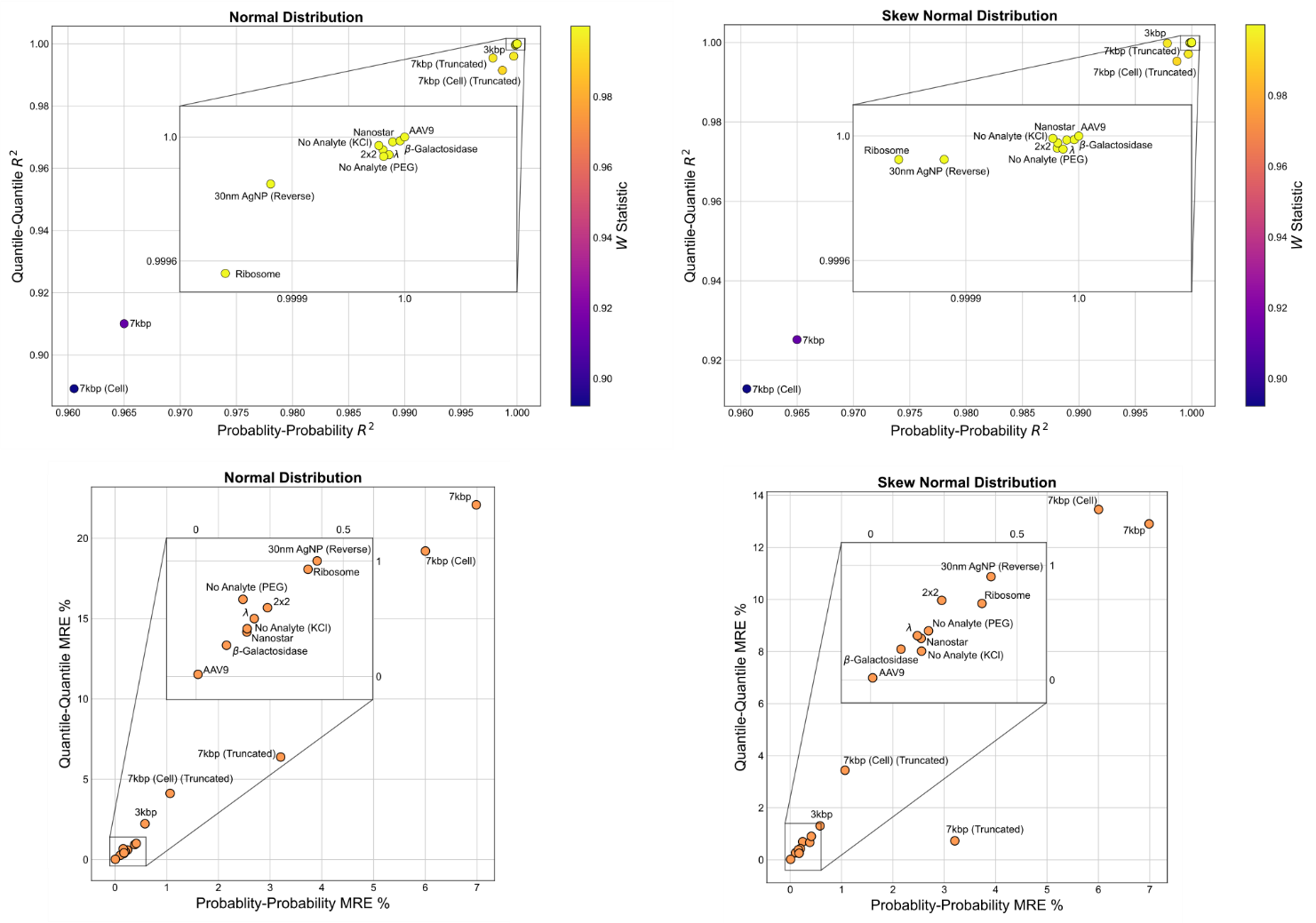
Fig. S6. Scatter plots of ensemble $\text{R}^{\boldsymbol{2}}$ and MRE results comparing trace distributions to normal and skew normal distributions. Methodology is provided in the Methods section of the main text.

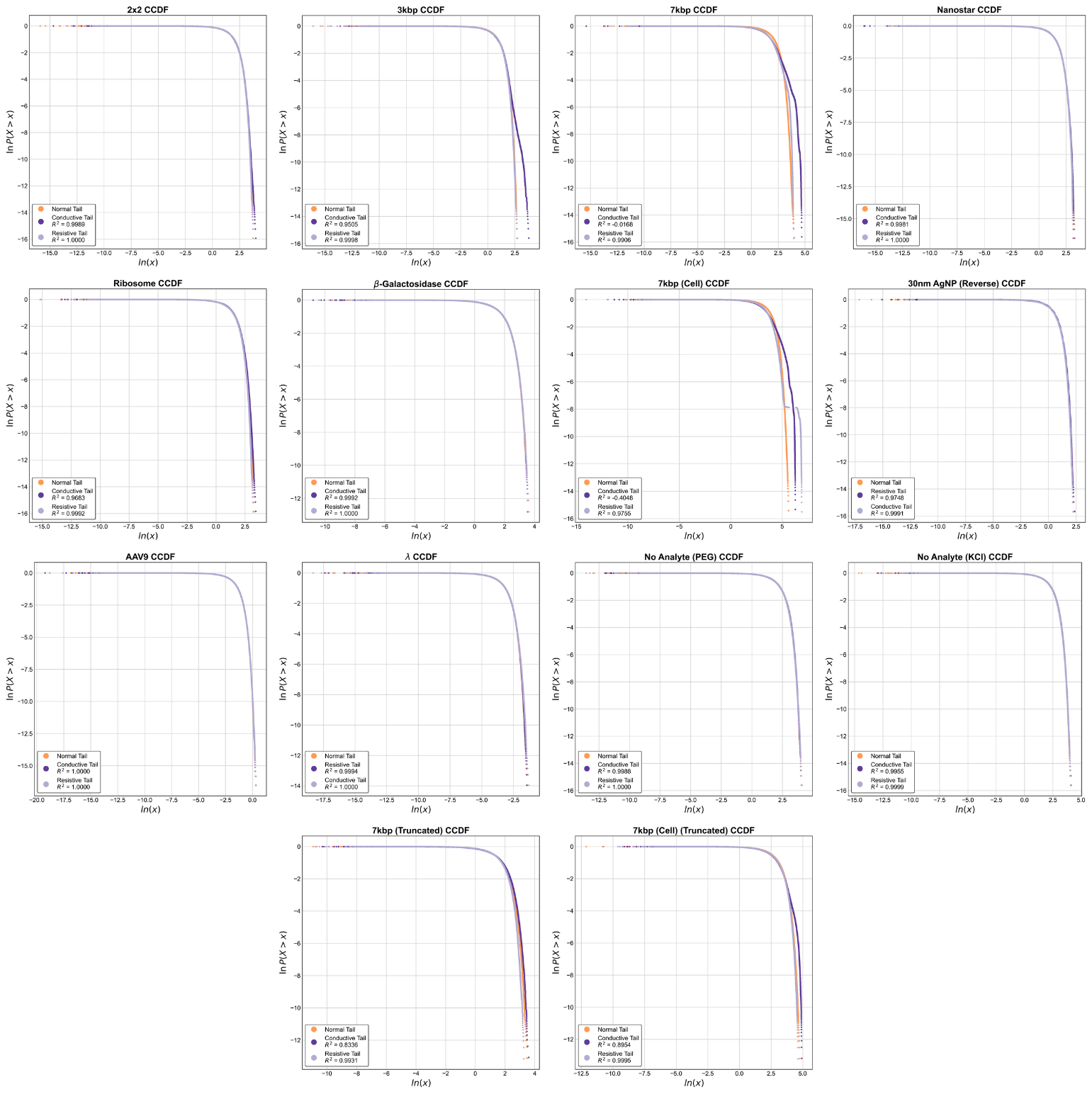
Fig. S7. CCDF plots comparing mean-split trace distributions with analogously split normal distributions. Methodology is provided in the Methods section of the main text.

| Process | R/S $\boldsymbol{H}$ | DFA $\boldsymbol{\alpha}$ | KPSS/ADF $\boldsymbol{I(d)}$ | PSD $\boldsymbol{\gamma}$ |
| --- | --- | --- | --- | --- |
| fGn | $0<H<0.5$ | $0<\alpha<0.5$ | $-0.5<d<0$ | $-1 < \gamma<0$ |
| wGn | $H=0.5$ | $\alpha=0.5$ | $d=0$ | $\gamma=0$ |
| fGn | $0.5<H<1$ | $0.5<\alpha<1$ | $0<d<0.5$ | $0 < \gamma<1$ |
| 1/f | $H = 1$ | $\alpha= 1$ | $d=0.5$ | $\gamma=1$ |
| fBm | $0<H<0.5$ | $1<\alpha<1.5$ | $0.5<d<1$ | $1 < \gamma<2$ |
| Bm | $H=0.5$ | $\alpha=1.5$ | $d=1$ | $\gamma= 2$ |
| fBm | $0.5<H<1$ | $1.5 <\alpha<2$ | $1 <d<1.5$ | $2 <\gamma<3$ |

**Table SII. Correspondence between Gaussian process classes and analysis output exponent ranges.**

| DFA $\boldsymbol{\alpha}$ | Self-Similarity | Dependence | Memory | Persistence | Temporal  Correlation | Marginal  Variance | Distributional Scaling |
| --- | --- | --- | --- | --- | --- | --- | --- |
| $\boldsymbol{0<\alpha<0.5}$ | *Second-order* | *Short-range*  *(stationary)* | *Short-term* | *Anti-persistent* | *Negative ACF (summable)* | *Constant* | *None* |
| $\boldsymbol{\alpha=0.5}$ | *None* | *None*  *(stationary)* | *None* | *Uncorrelated* | *Zero ACF* | *Constant* | *None* |
| $\boldsymbol{0.5<\alpha<1}$ | *Second-order* | *Long-range*  *(stationary)* | *Long-term* | *Persistent* | *Positive ACF*  *(non-summable)* | *Constant* | *None* |
| $\boldsymbol{\alpha= 1}$ | *Critical*  *boundary* | *Critical*  *boundary* | *Critical*  *boundary* | *Perfect persistence*  *limit* | *Constant ACF*  *(non-decaying)* | *Logarithmic*  *divergence* | *Critical*  *boundary* |
| $\boldsymbol{1<\alpha<1.5}$ | *Exact* | *Long-range*  *(non-stationary,*  *structural)* | *Increment*  *short-term* | *Increment*  *anti-persistent* | *Increment*  *negative ACF* | $\propto t^{2\alpha-2}$  *(subdiffusive)* | $X\left( at \right)\overset{d}{=}a^{\alpha-1}X\left( t \right)$ |
| $\boldsymbol{\alpha=1.5}$ | *Exact* | *Long-range*  *(non-stationary,*  *structural)* | *Increment*  *uncorrelated* | *Increment*  *uncorrelated* | *Increment*  *zero ACF* | $\propto t$  *(diffusive)* | $X\left( at \right)\overset{d}{=}a^{0.5}X\left( t \right)$ |
| $\boldsymbol{1.5<\alpha<2}$ | *Exact* | *Long-range*  *(non-stationary,*  *structural)* | *Increment*  *long-term* | *Increment*  *Persistent* | *Increment*  *positive ACF* | $\propto t^{2\alpha-2}$  *(superdiffusive)* | $X\left( at \right)\overset{d}{=}a^{\alpha-1}X\left( t \right)$ |

**Table SIII.** **Summary of the interpretation of DFA exponent** $\boldsymbol{\alpha}$ **over its conventional range, showing the associated properties of the resulting Gaussian process.**

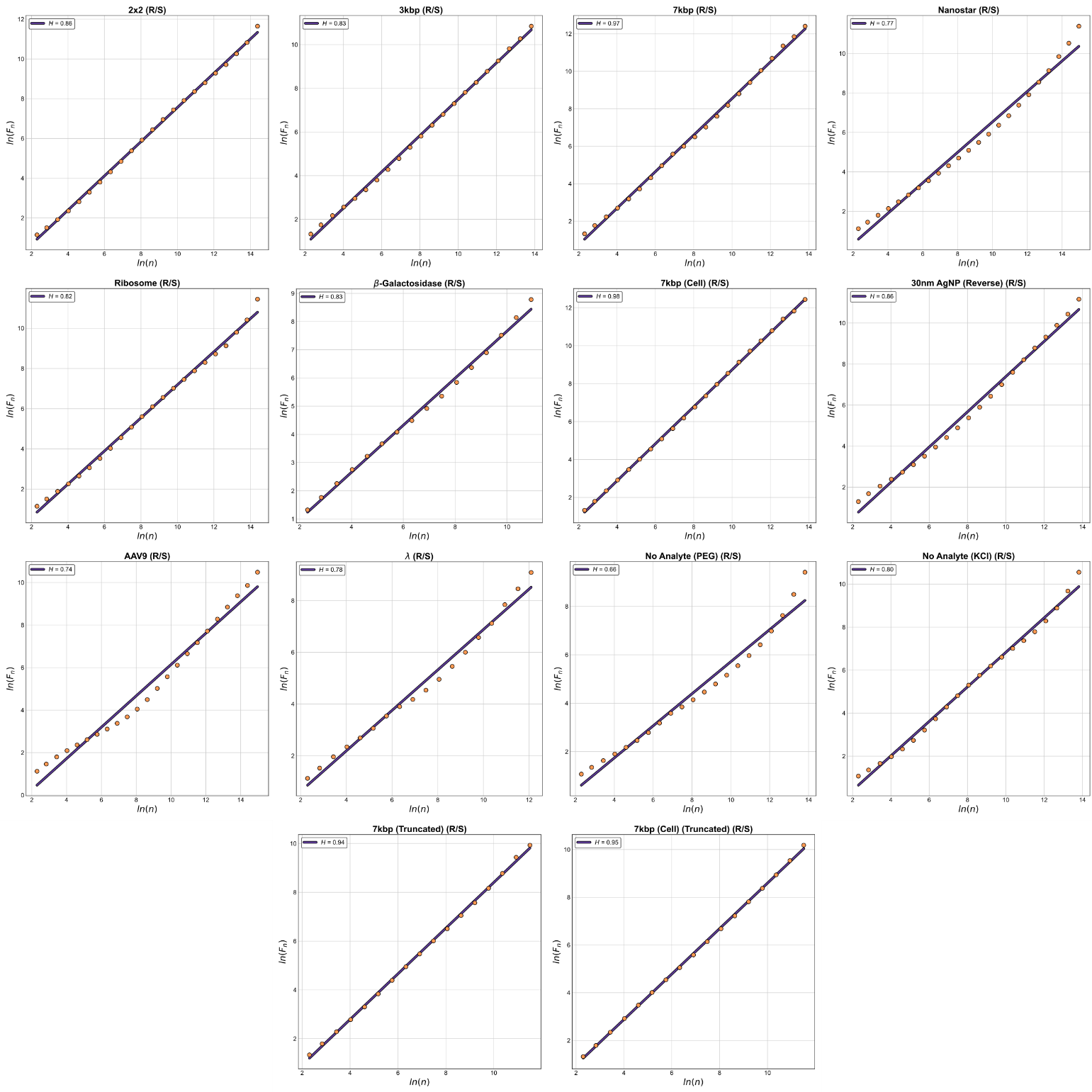
Fig. S8. R/S plots with linear fits for the fourteen traces. Methodology is provided in the Methods section of the main text.

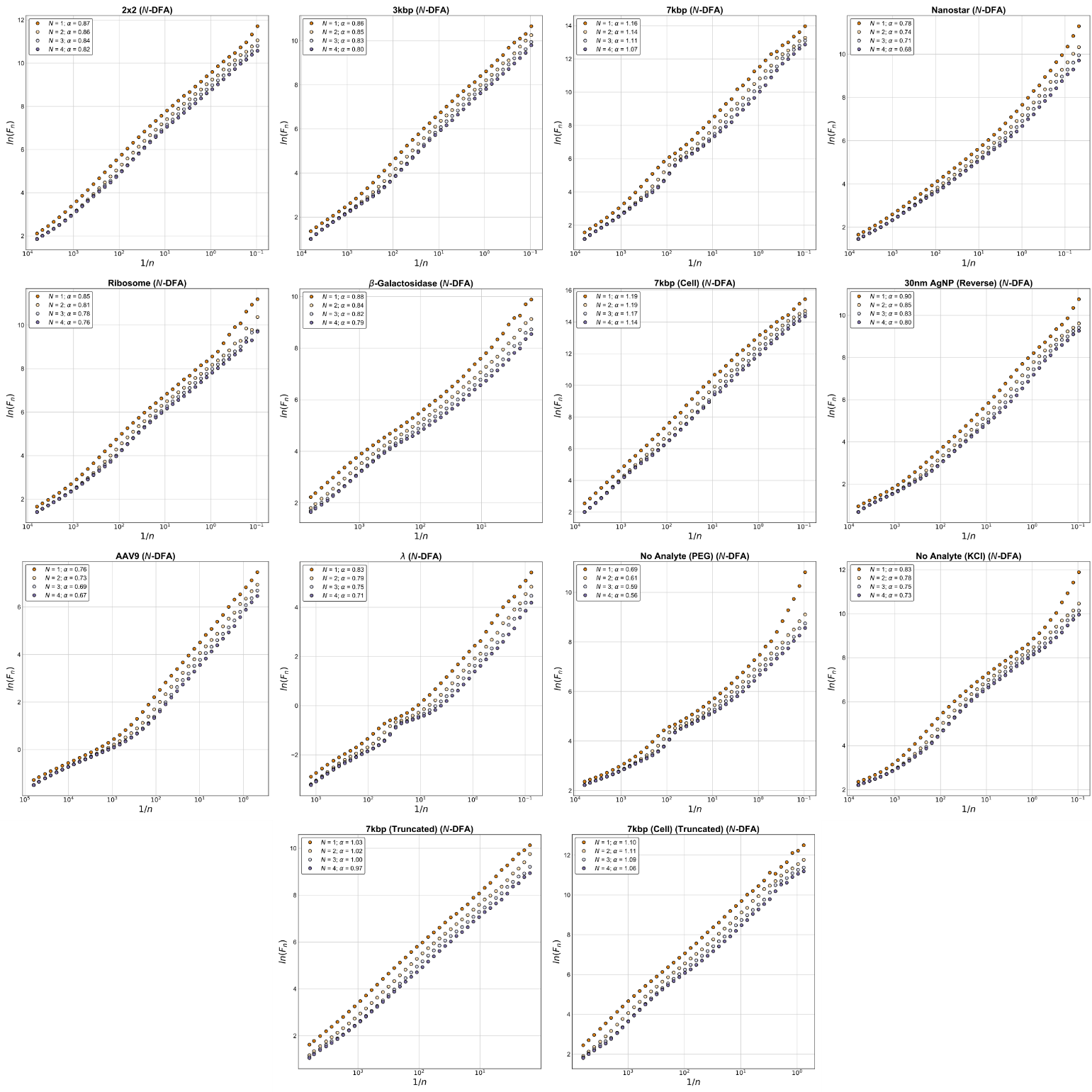
Fig. S9. N-DFA plots with linear fits for the fourteen traces. Methodology is provided in the Methods section of the main text.

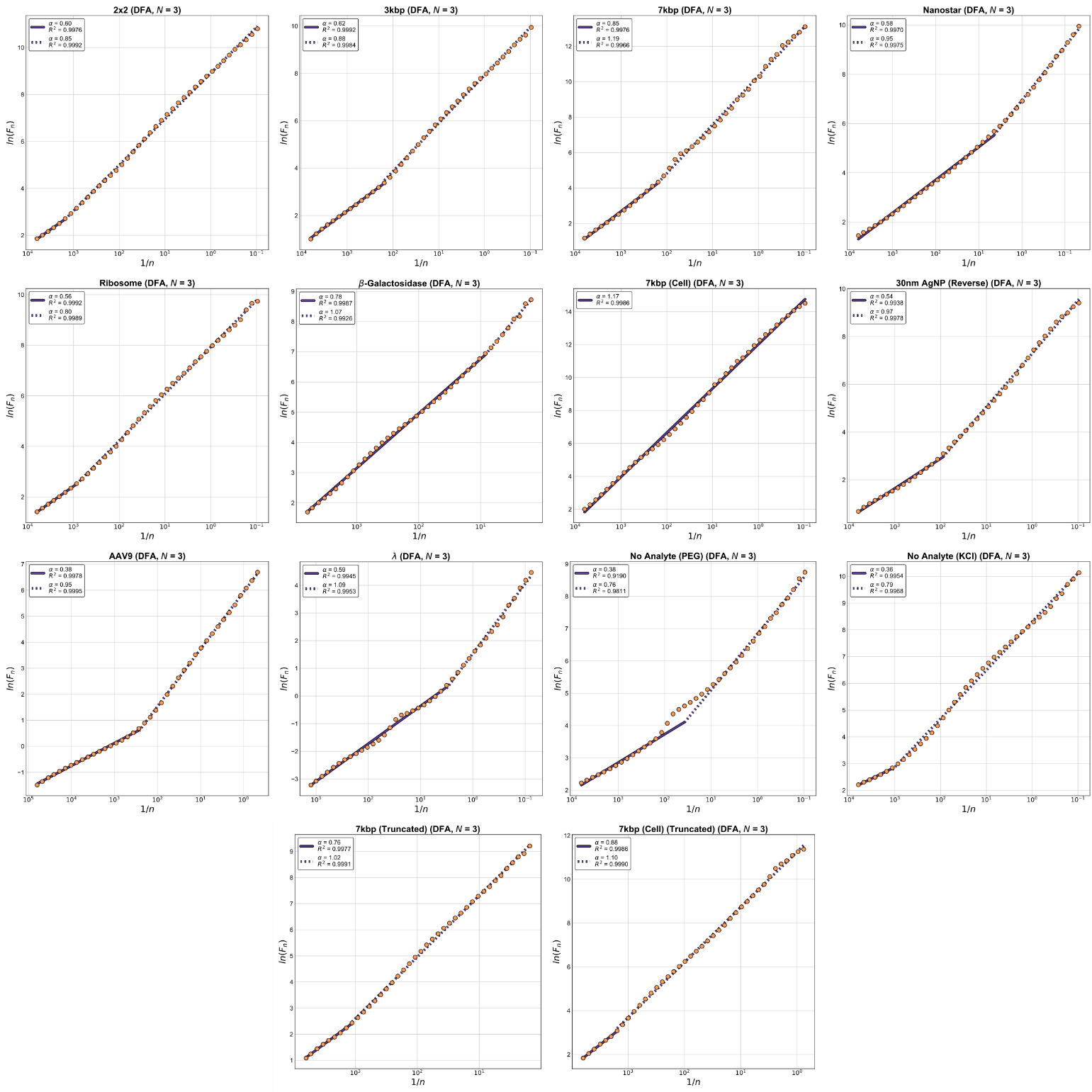
Fig. S10. Final DFA plots ($\boldsymbol{N}$= 3) with piecewise fits for the fourteen traces. Methodology is provided in the Methods section of the main text.

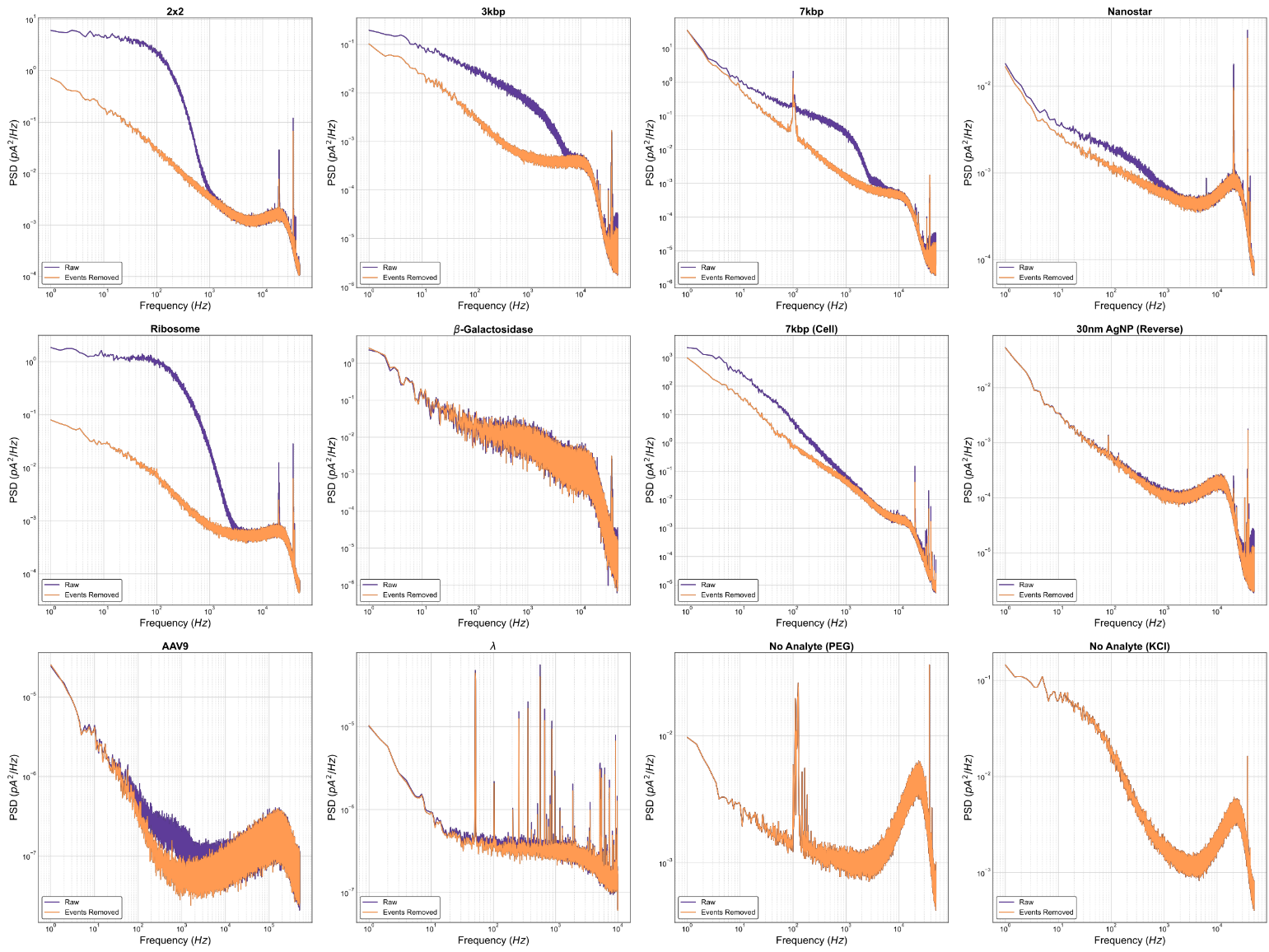
Fig. S11. PSD plots for the twelve traces with and without events. Methodology is provided in the Methods section of the main text.
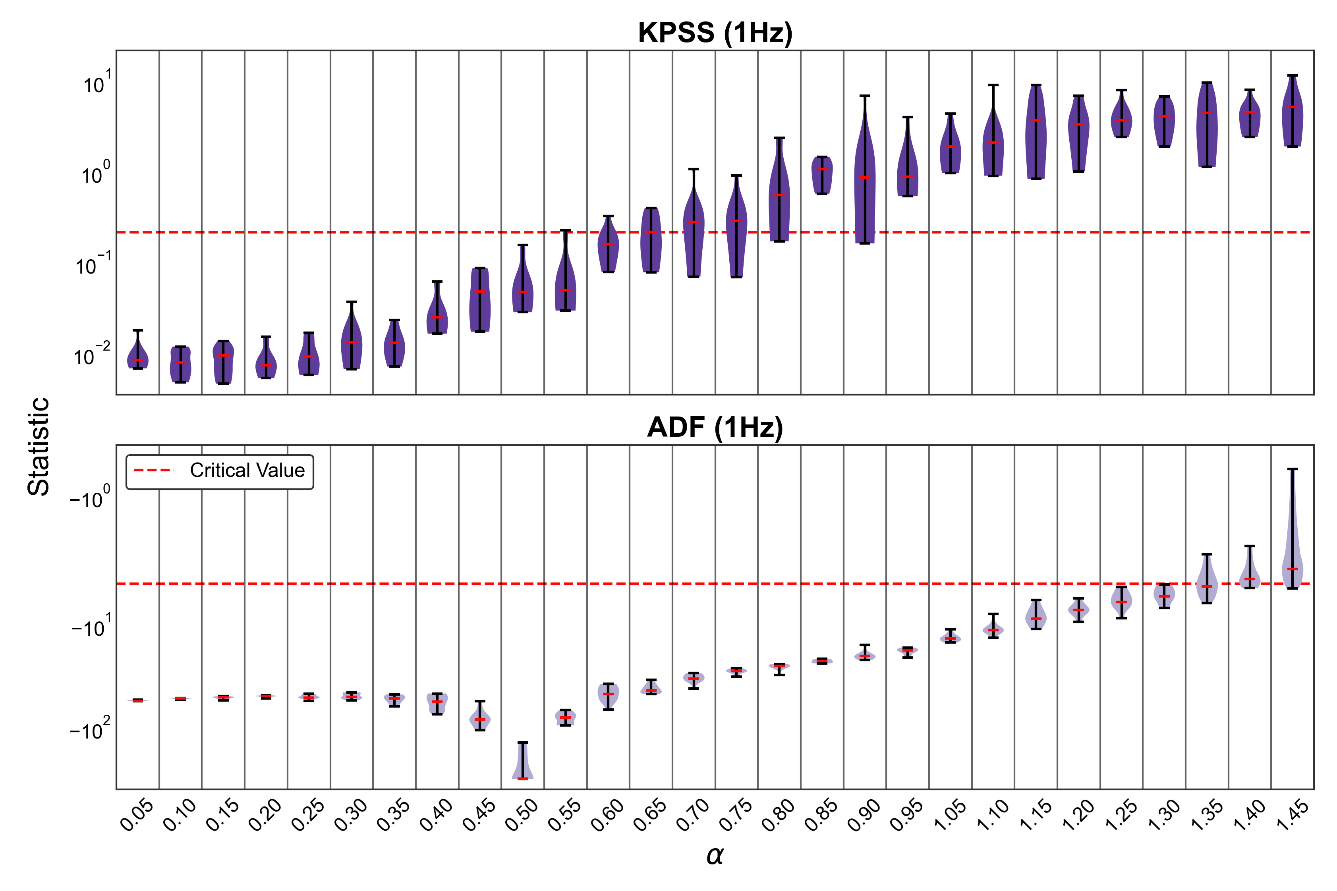

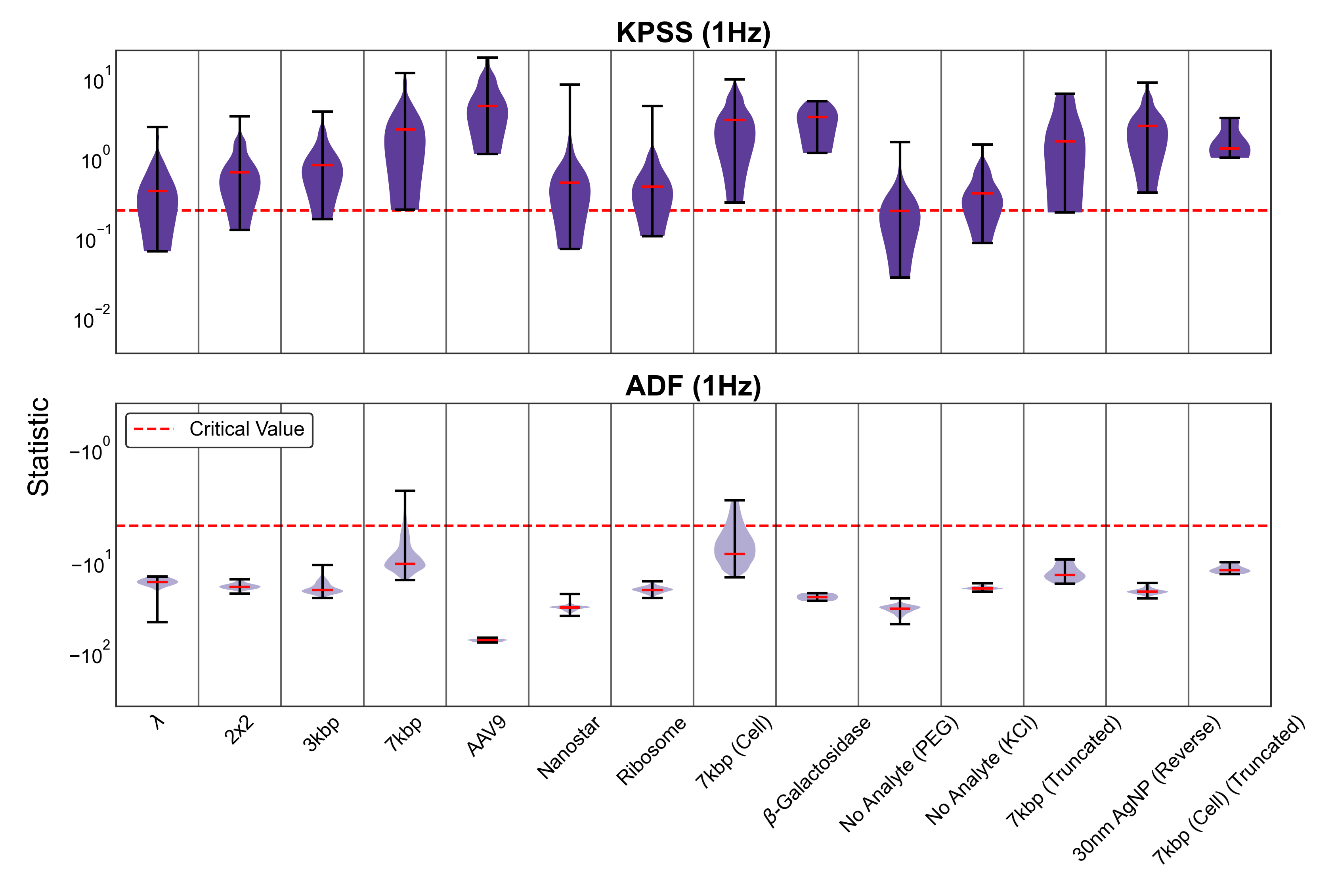

Fig. S12. KPSS and ADF statistic distributions for the reference set and the fourteen traces.

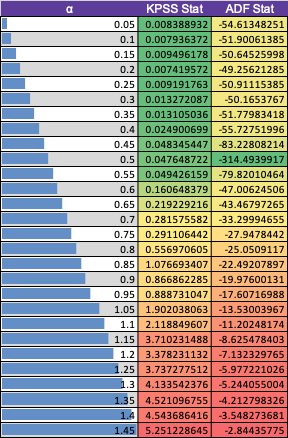

Table SIV. KPSS and ADF statistics for the reference set.

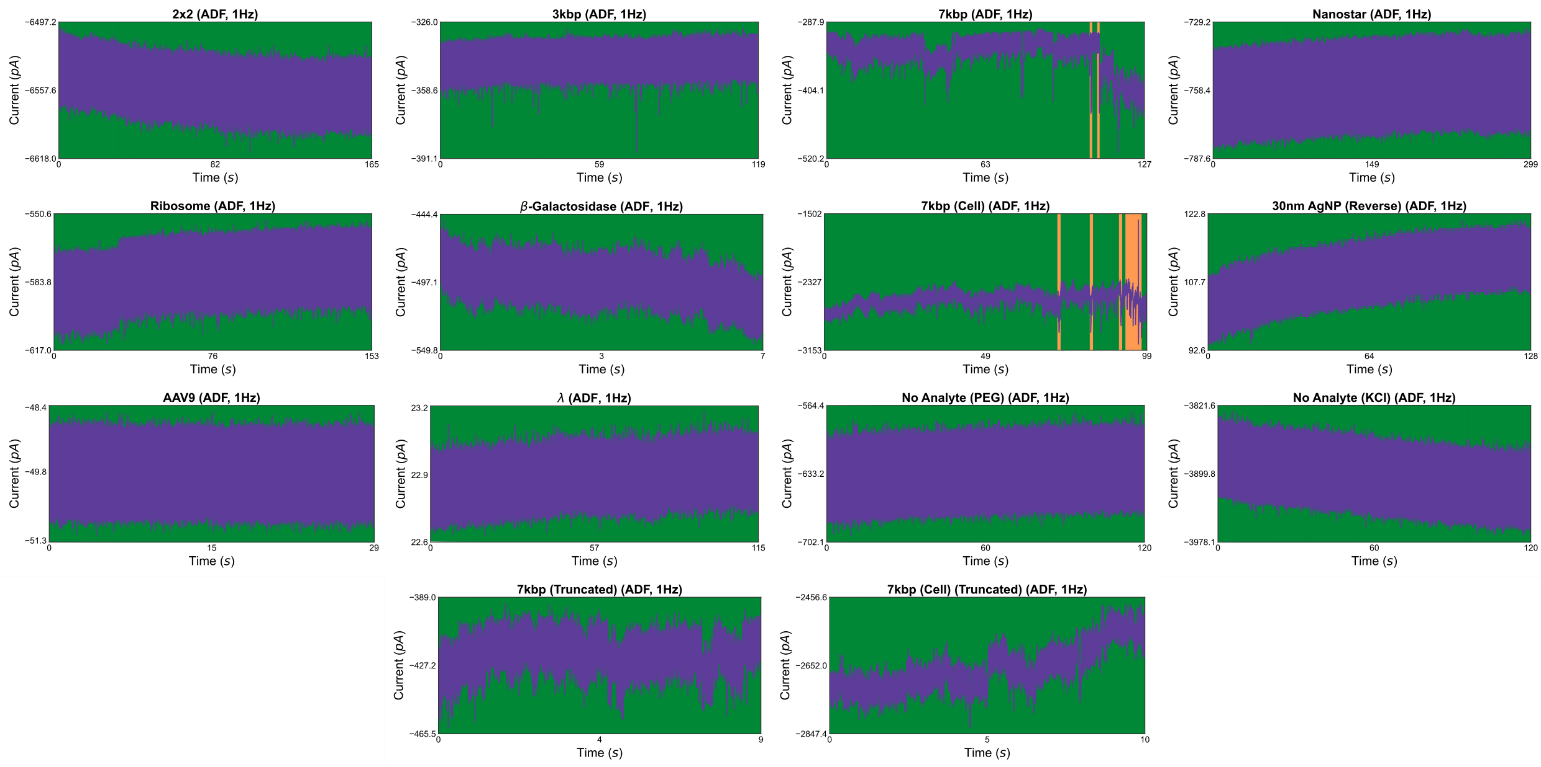
Fig. S13. ADF assertion and rejection plots for the fourteen traces. Methodology is provided in the Methods section of the main text.

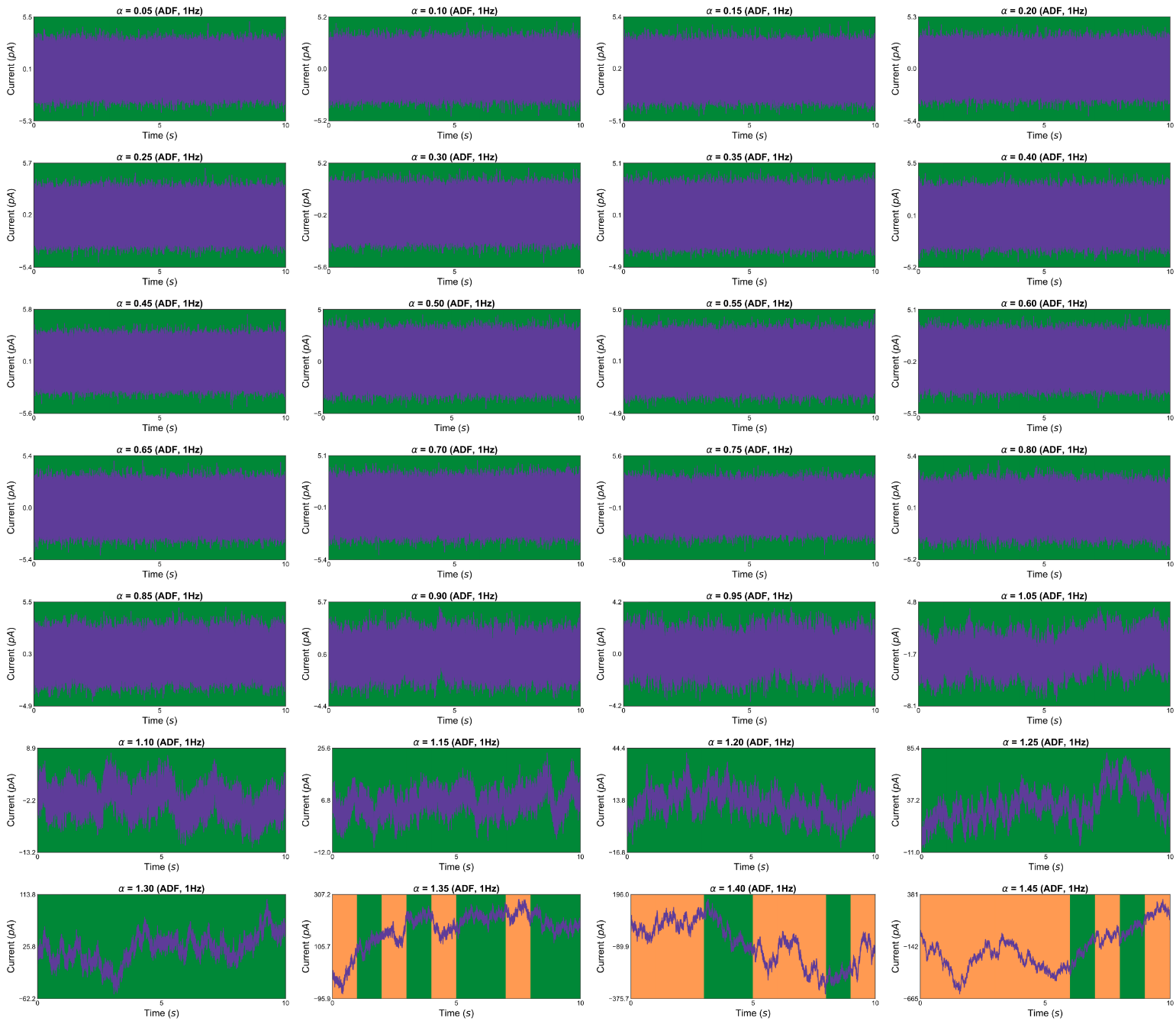
Fig. S14. ADF assertion and rejection plots for the reference set. Methodology is provided in the Methods section of the main text.

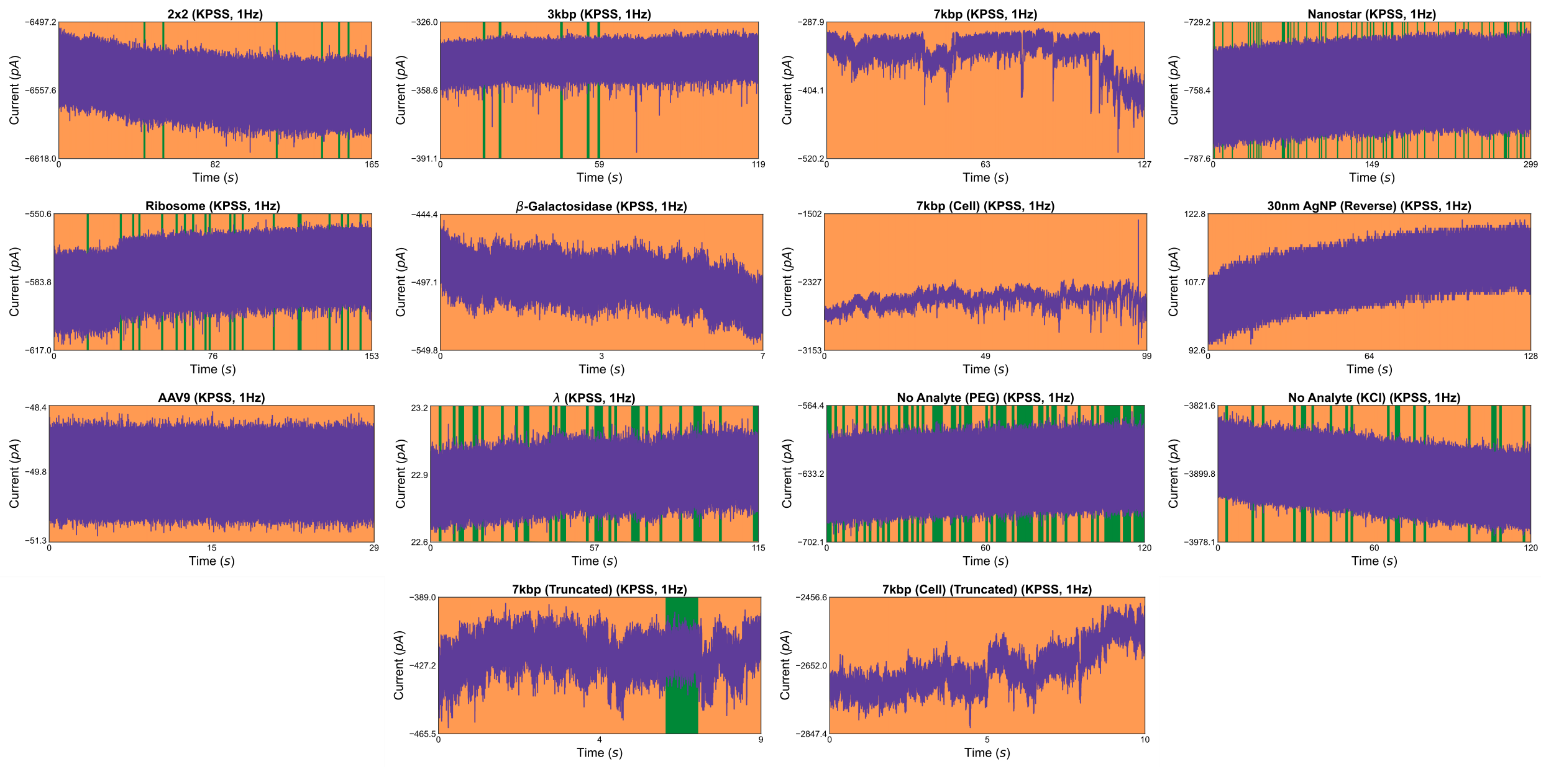
Fig. S15. KPSS assertion and rejection plots for the fourteen traces. Methodology is provided in the Methods section of the main text.

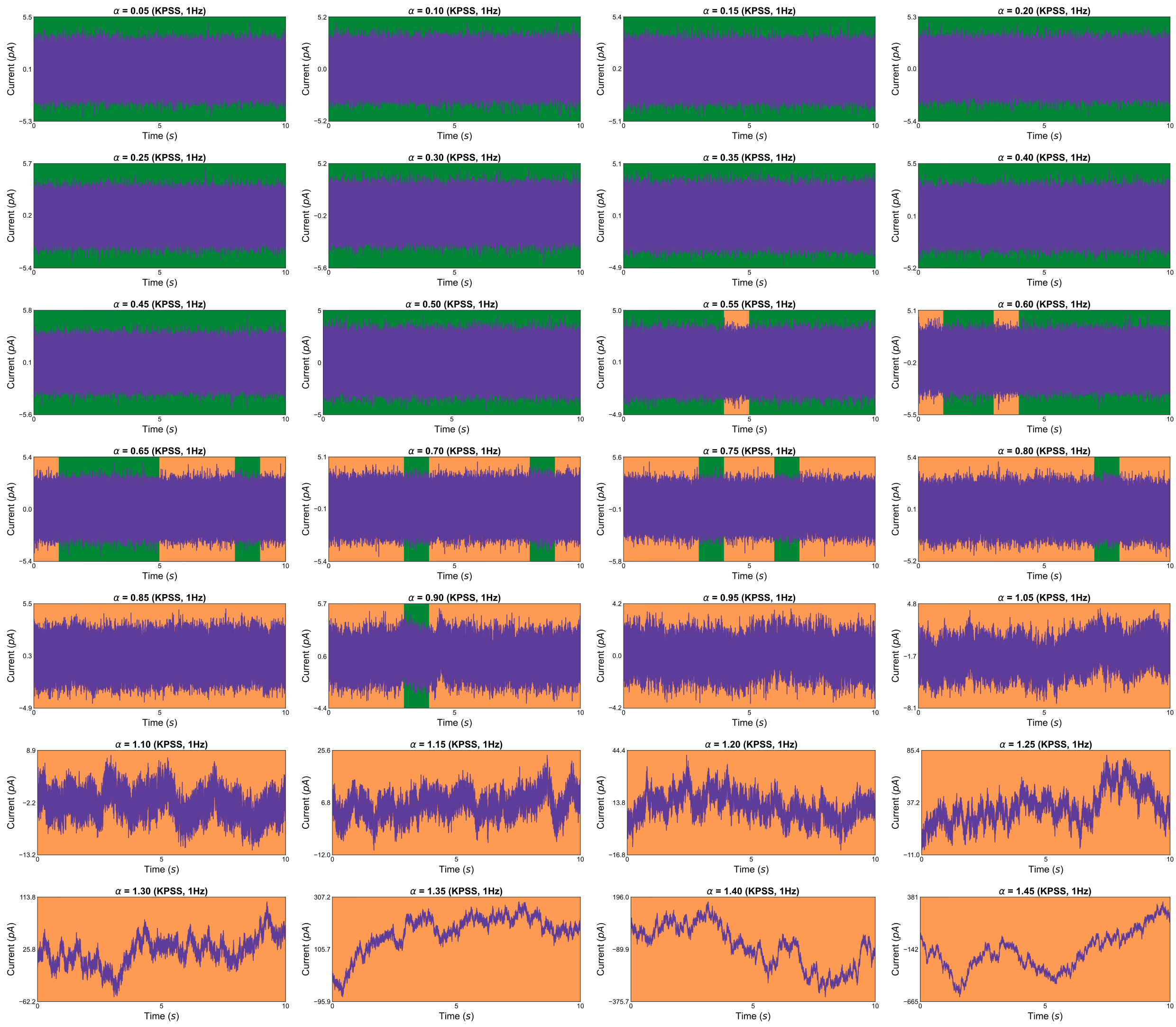
Fig. S16. KPSS assertion and rejection plots for the reference set. Methodology is provided in the Methods section of the main text.

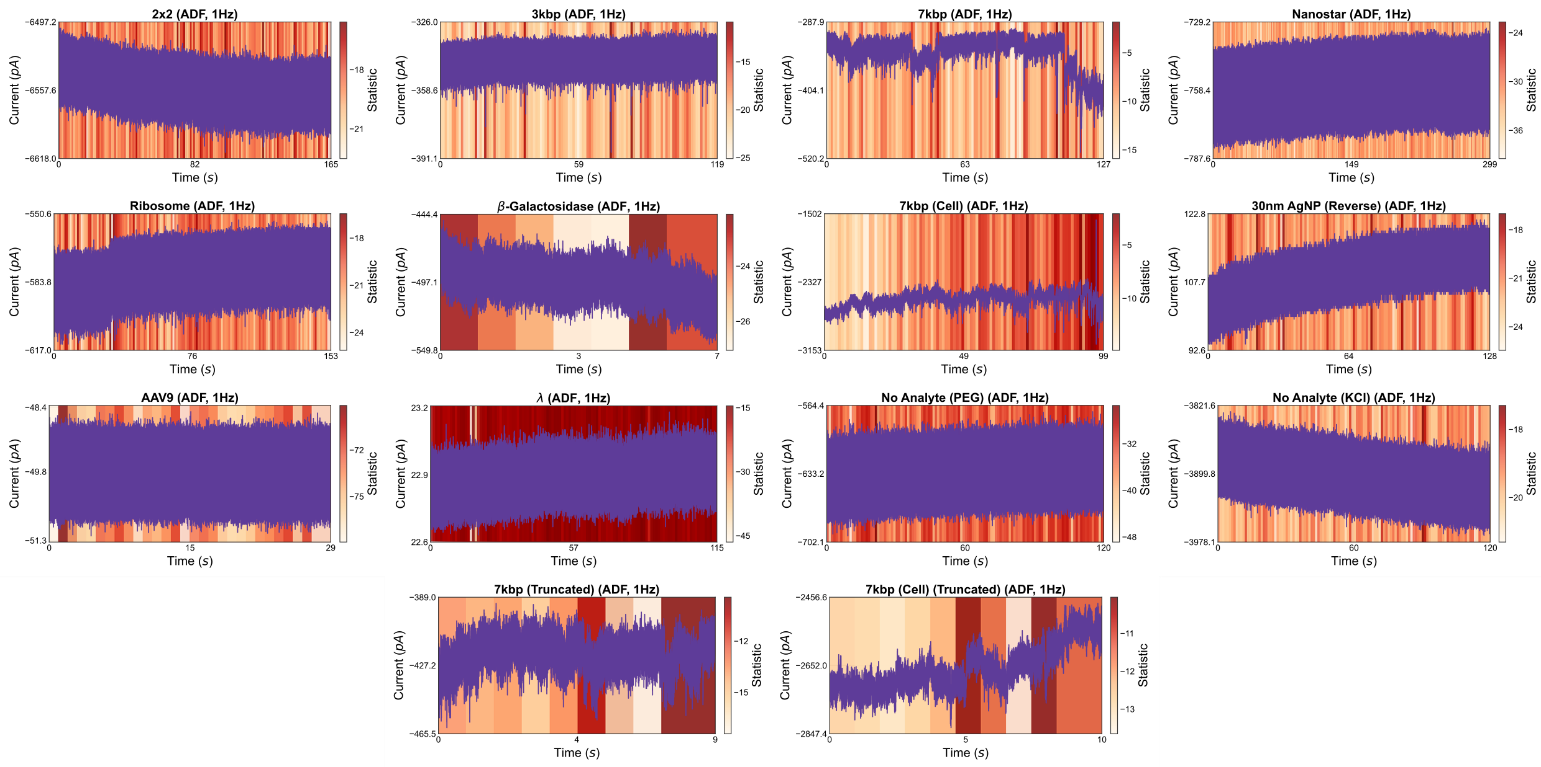
Fig. S17. ADF statistic plots for the fourteen traces. Methodology is provided in the Methods section of the main text.

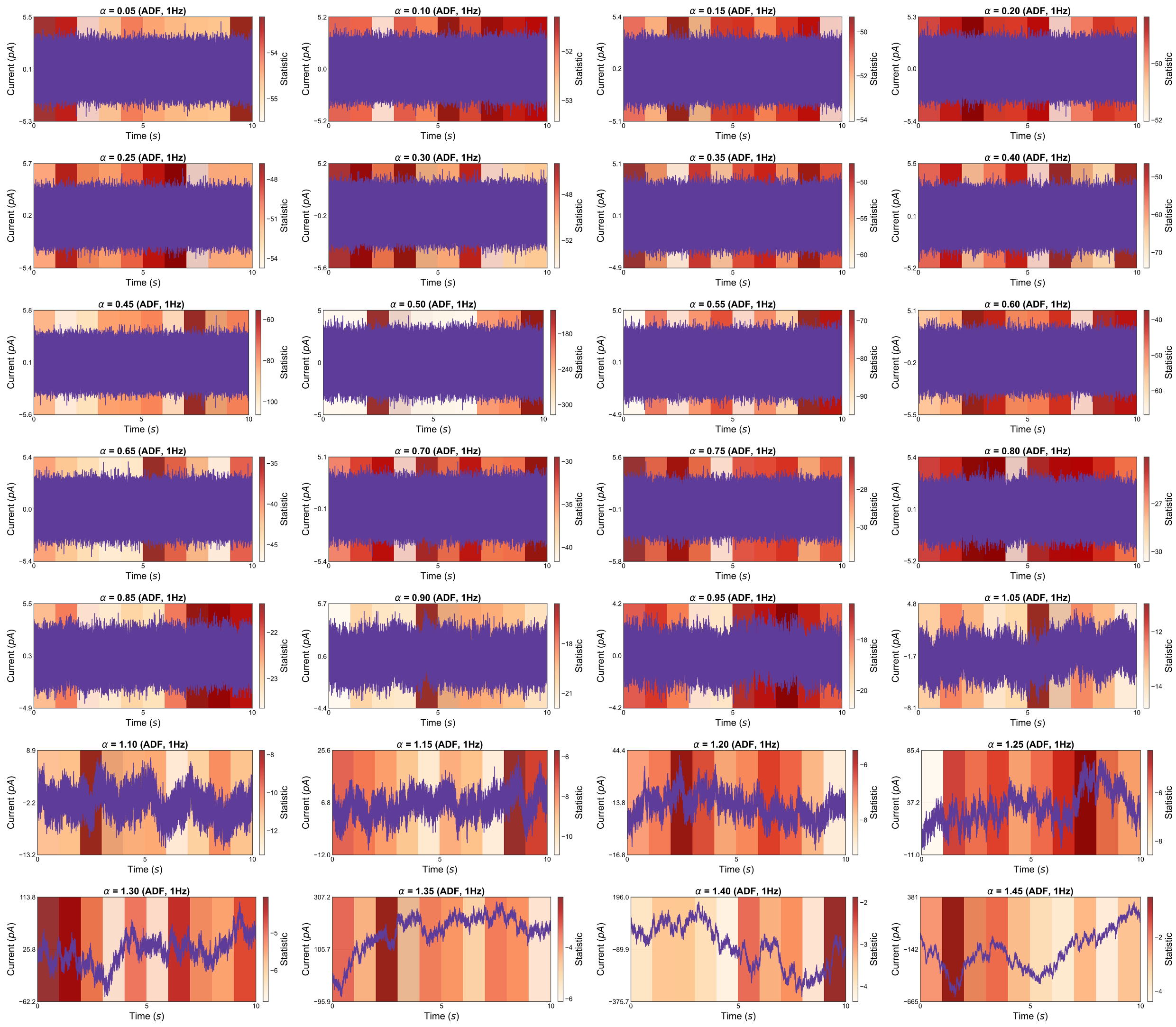
Fig. S18. ADF statistic plots for the reference set. Methodology is provided in the Methods section of the main text.

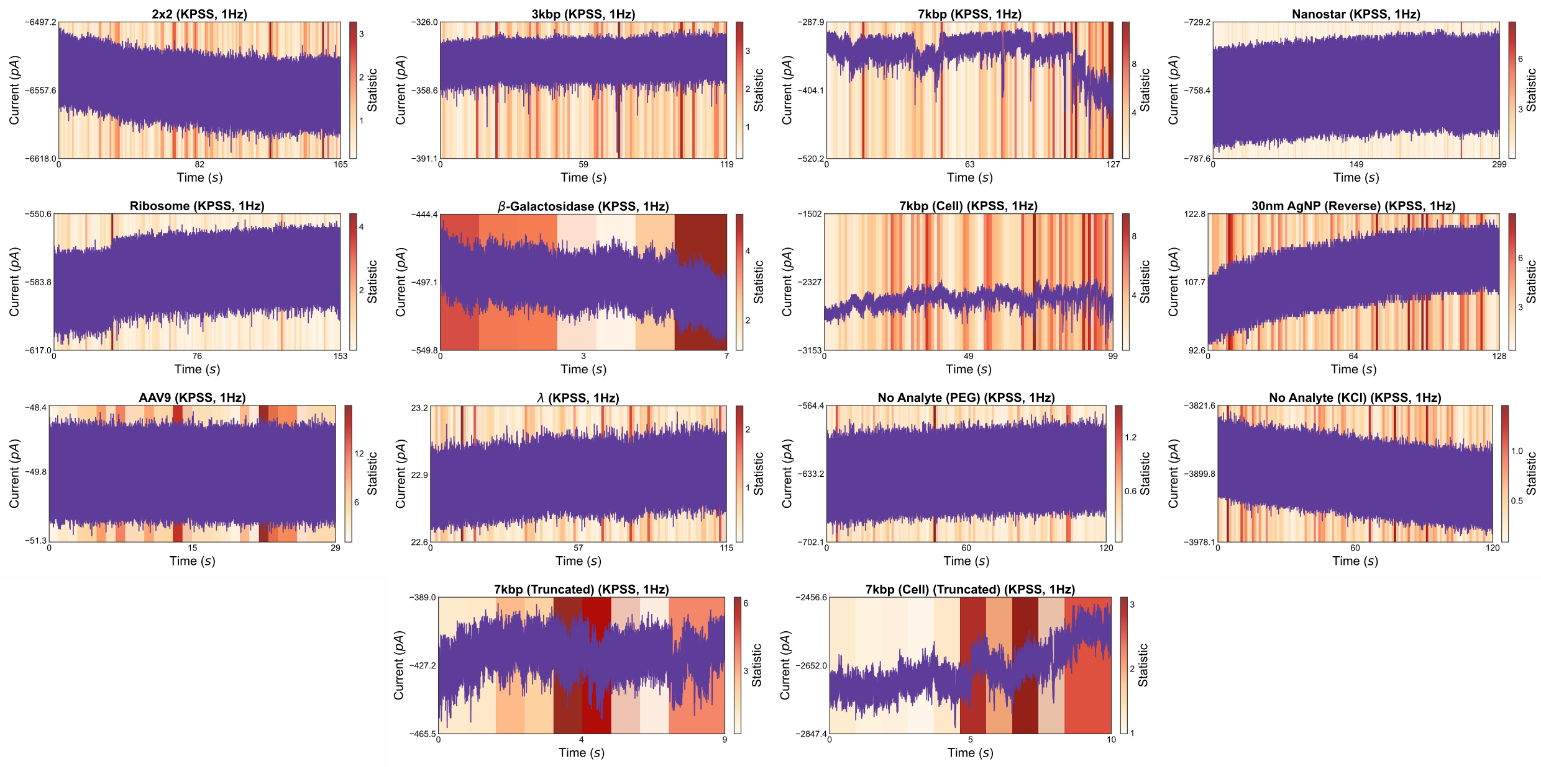
Fig. S19. KPSS statistic plots for the fourteen traces. Methodology is provided in the Methods section of the main text.

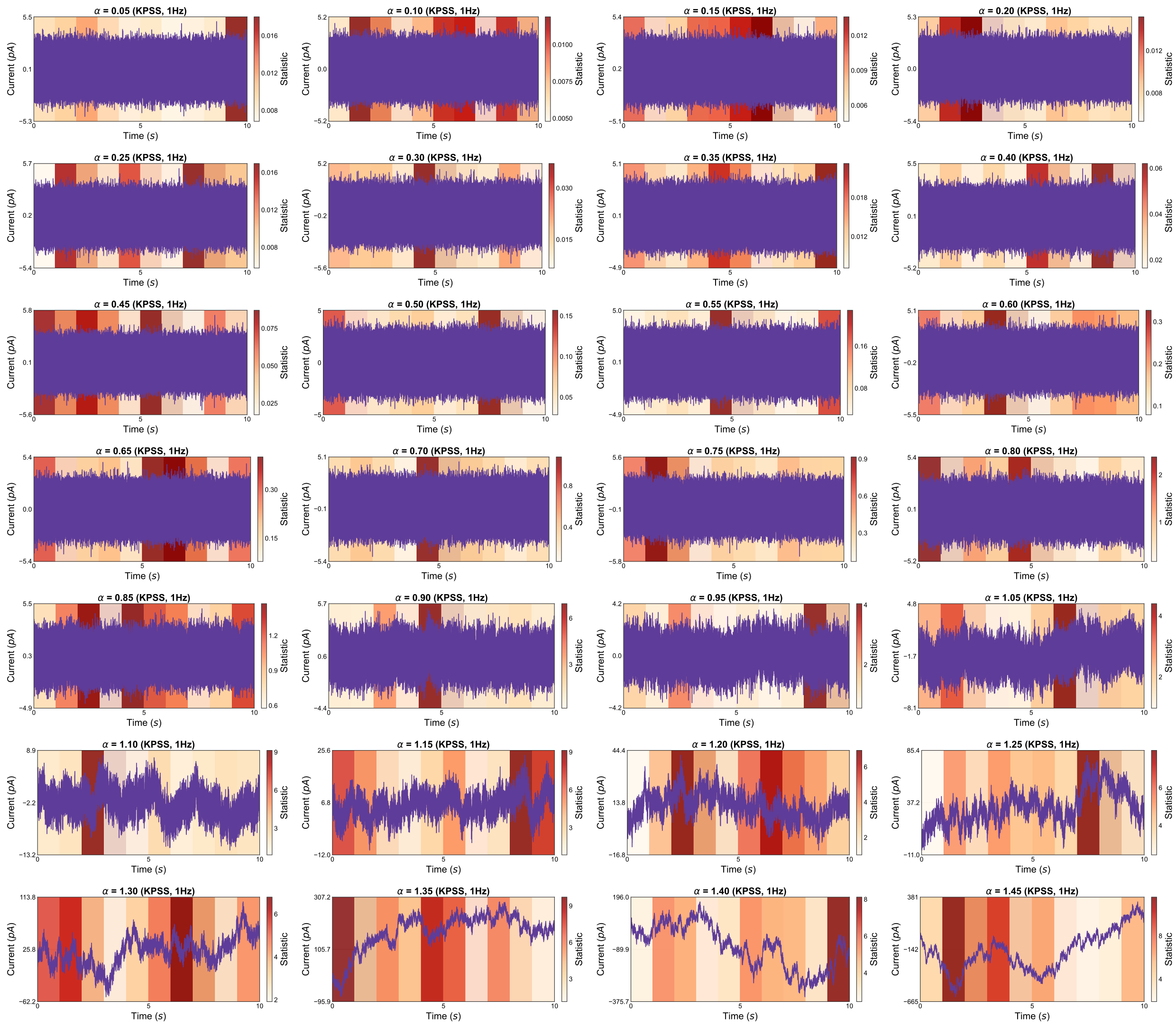
Fig. S20. KPSS statistic plots for the reference set. Methodology is provided in the Methods section of the main text.

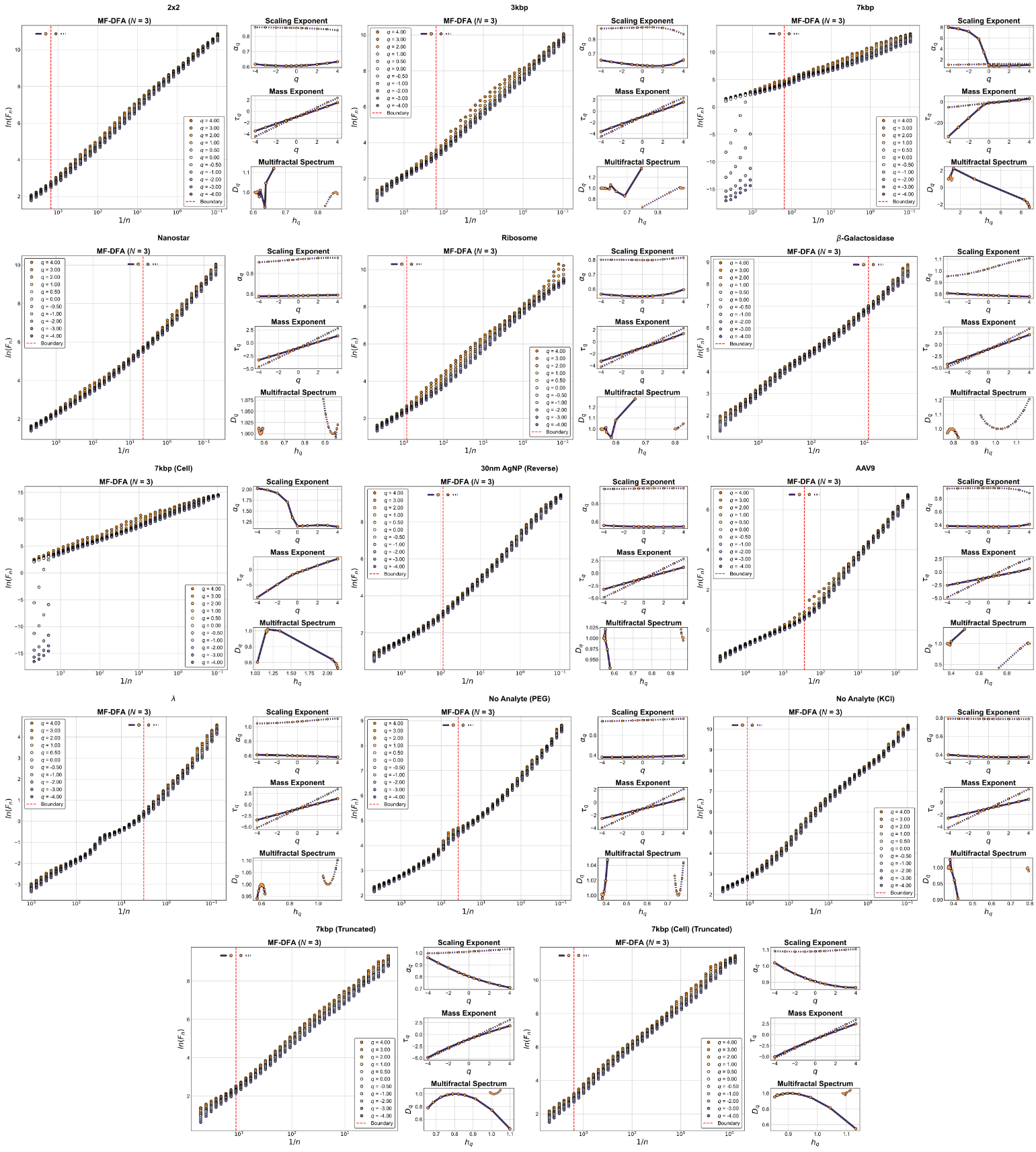
Fig. S21. MF-DFA plots ($\boldsymbol{N}$= 3) for the fourteen traces. Methodology is provided in the Methods section of the main text.

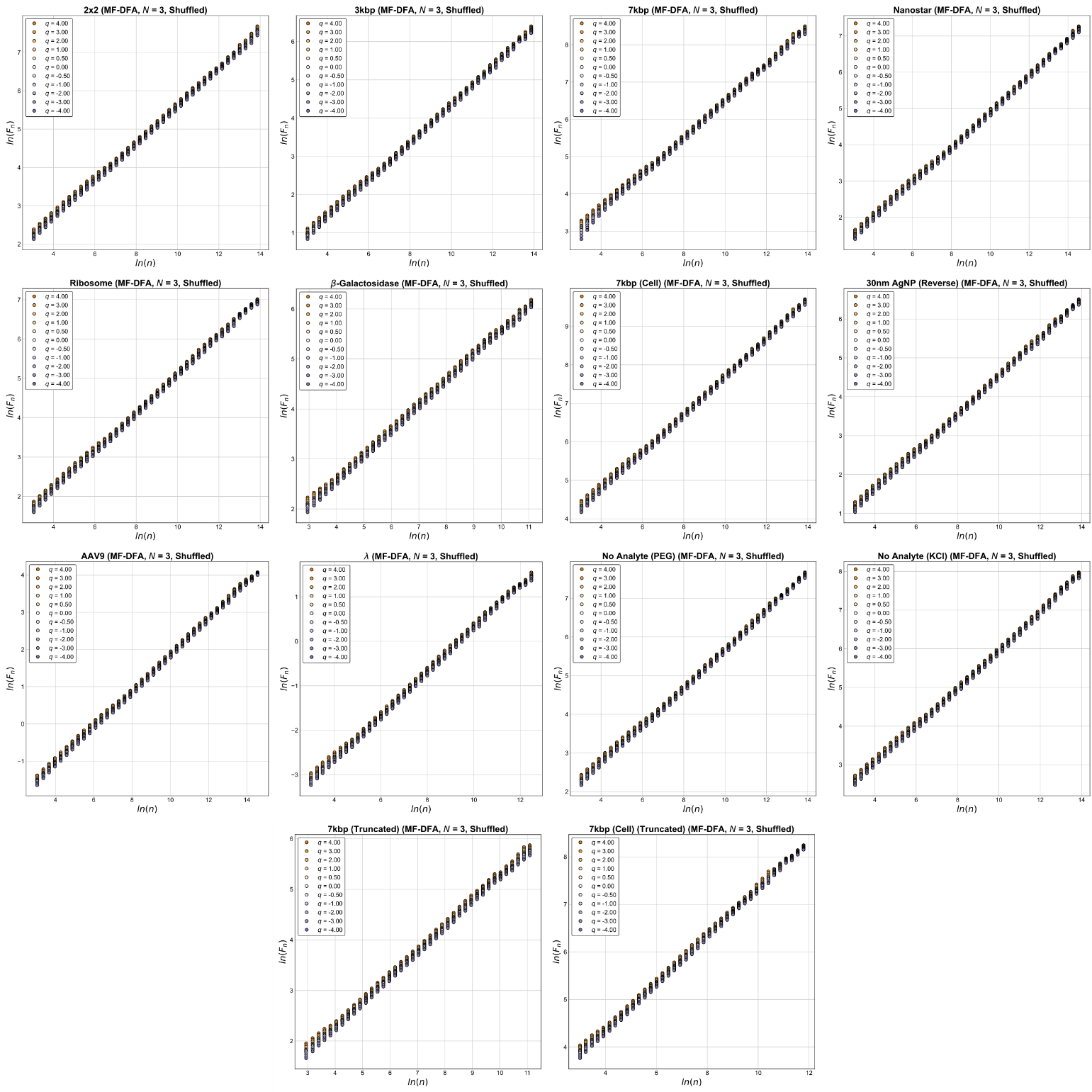
Fig. S22. Shuffled MF-DFA plots ($\boldsymbol{N}$= 3) for the fourteen traces. Methodology is provided in the Methods section of the main text.

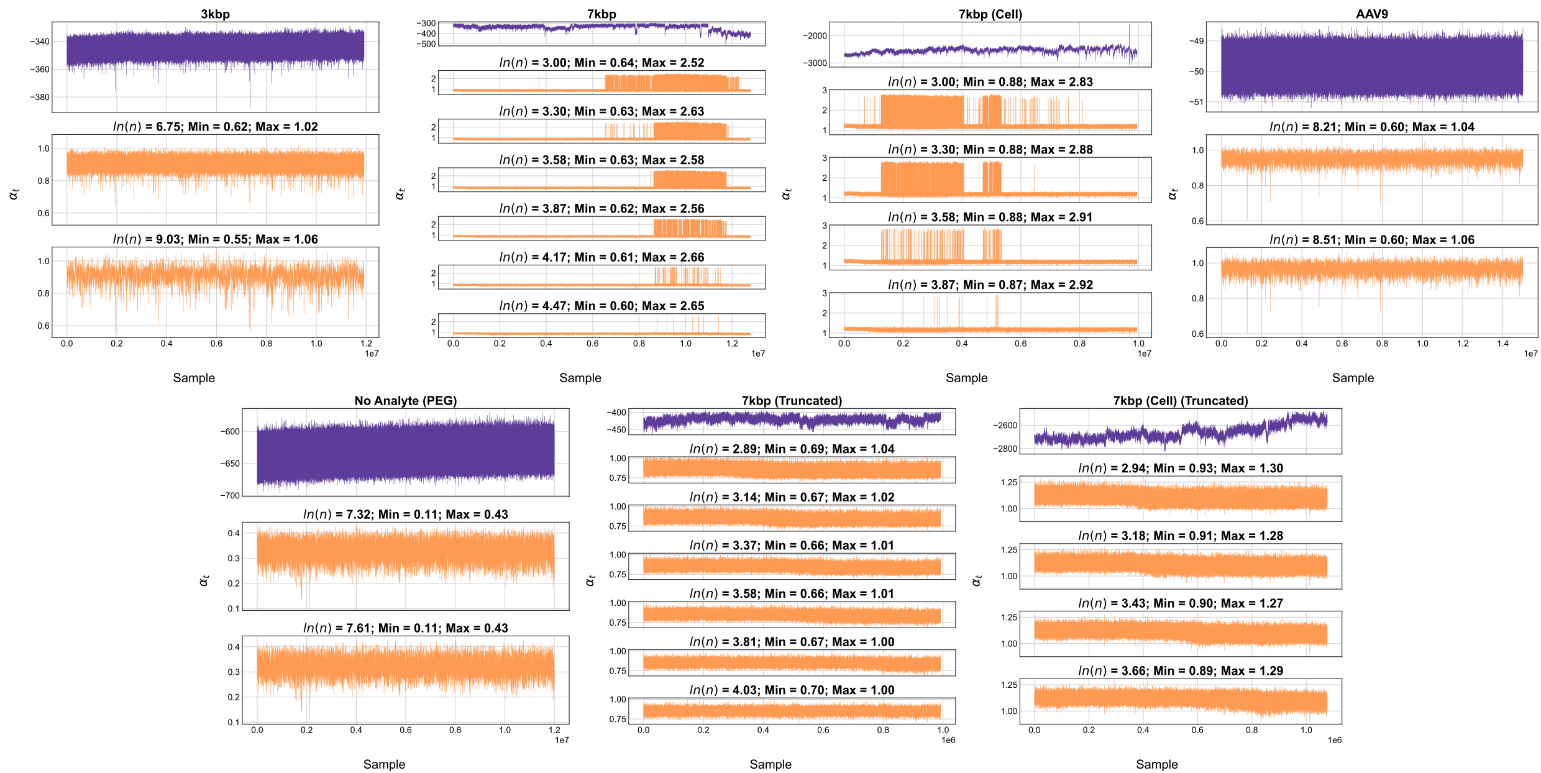
Fig. S23. T-DFA plots for the traces exhibiting apparent multifractality. Methodology is provided in the Methods section of the main text.

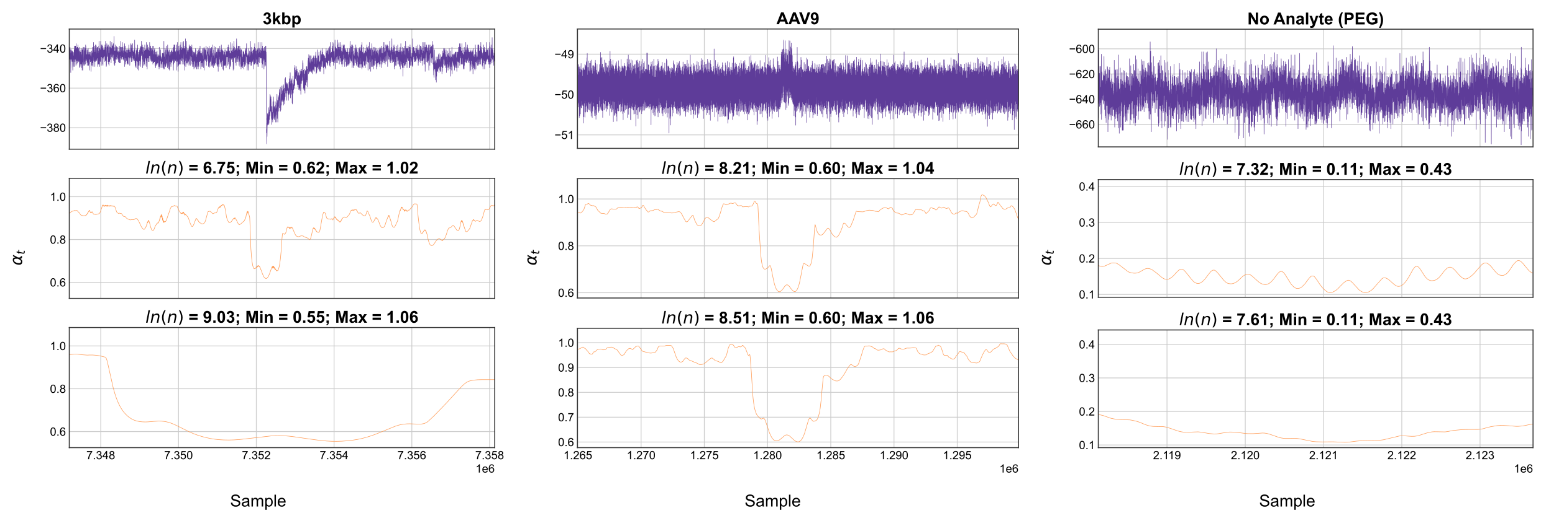
Fig. S24. Zoomed in T-DFA plots for the traces exhibiting apparent multifractality. Methodology is provided in the Methods section of the main text.

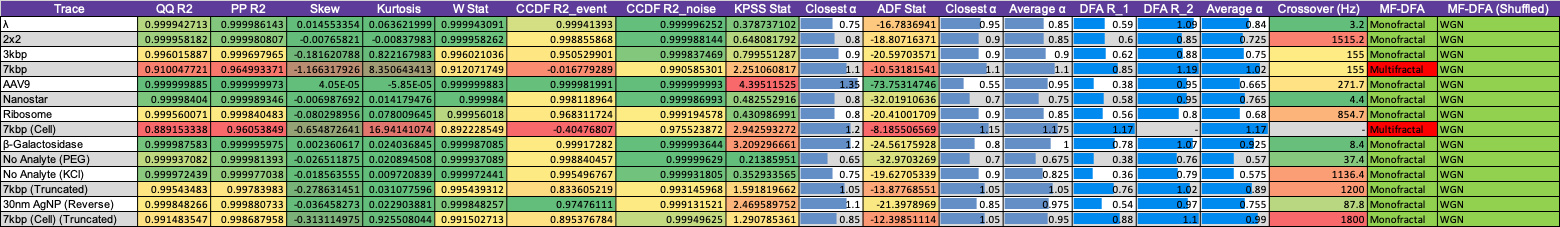
Table SV. Overall results for the fourteen traces.

### Practical Analysis Pipeline for Trace Validation

Here we describe a condensed and practical version of the analysis procedure presented in the main text so researchers can validate their own data against the proposed model. This procedure is focused on the most important aspects of the analyses. Complete methodology and interpretation guidelines are found in the main text Sec. II, and information on method synthesis is available in main text Sec. III.

1. Remove events from the trace using an appropriate algorithm and concatenate the remaining segments. In this manuscript, we adopt a modified iterative baseline method with a secondary manual pass for improved accuracy (main text Sec. II B). However, alternative algorithms may be preferable depending on the trace, and the second pass is non-essential.
2. Conduct a visual analysis to identify prominent trace features which may bias the following analyses. If structural breaks are identified, truncate the trace to a stable span, focusing on maximising the remaining sample size.
3. Prior to distributional analyses, identify the observation time quasi-deterministic trend and apply nonlinear detrending using a sensible equivalent cutoff frequency. Here, we found 1 Hz to be appropriate for our data, but this may be insufficient for other traces.
4. For distributional analysis, compute quantile-quantile (Q-Q) and probability-probability (P-P) plots against normal distributions for the detrended trace following the procedure in main text Sec. II D. If structural breaks were previously identified, comparison against a skew-normal distribution may be preferable. If major divergences are present, complementary cumulative distribution functions (CCFFs) will provide additional insight, and should be computed as described in main text Sec. II E. Intrinsic divergence from normality will be identifiable through these three analyses, and artefactual divergences will be clear in the tail-specific CCDFs combined with prior visual analyses.
5. Second-order and multifractal analyses should be combined via Multifractal Detrended Fluctuation Analysis (MF-DFA) using the method and interpretation described in main text Sec. II. K, where second-order exponents can be deduced from the $q = 2$ result. Power Spectral Density (PSD) and Scaled Stationarity analyses should be considered if divergences are present in the MF-DFA result, enabling confirmation of periodic interference and progressive non-stationarities, respectively. Moreover, Time-dependent Detrended Fluctuation Analysis (T-DFA) (main text Sec. II. L) can be used to on divergent scales to classify time-localised divergences.
6. Finally, MF-DFA should be computed on the shuffled trace to complete the interpretation of the prior steps (main text Sec. II. K1). Uncorrelated white noise scaling enables interpretation of the $q = 2$ unshuffled result regimes as fractional Gaussian noise (fGn) or fractional Brownian motion (fBm) exponents, contingent on prior identification of approximate normality and absence of intrinsic multifractality. Otherwise, this result will provide corroboration for previously identified divergences related to the interpretation in main text Sec. II. K1.

If the trace does not adhere to the model, identification of an alternative characterisation is dependent on the specific unworthy descriptors. For example, alternative distributions can be tested against, but modifications to the following analyses will have to be made as either their interpretation or assumptions are based on normality. Variations of these tests for alternative distributions can be found in the literature.
